# MEDICC2: whole-genome doubling aware copy-number phylogenies for cancer evolution

**DOI:** 10.1101/2021.02.28.433227

**Authors:** Tom L Kaufmann, Marina Petkovic, Thomas BK Watkins, Emma C Colliver, Sofya Laskina, Nisha Thapa, Darlan C Minussi, Nicholas Navin, Charles Swanton, Peter Van Loo, Kerstin Haase, Maxime Tarabichi, Roland F Schwarz

## Abstract

Chromosomal instability (CIN) and somatic copy-number alterations (SCNA) play a key role in the evolutionary process that shapes cancer genomes. SC-NAs comprise many classes of clinically relevant events, such as localised amplifications, gains, losses, loss-of-heterozygosity (LOH) events, and recently discovered parallel evolutionary events revealed by multi-sample phasing. These events frequently appear jointly with whole genome doubling (WGD), a transformative event in tumour evolution involving tetraploidization of genomes preceded or followed by individual chromosomal copy-number changes and associated with an overall increase in structural CIN.

While SCNAs have been leveraged for phylogeny reconstruction in the past, existing methods do not take WGD events into account and cannot model parallel evolution. They frequently make use of the infinite sites assumption, do not model horizontal dependencies between adjacent genomic loci and can not infer ancestral genomes. Here we present MEDICC2, a new phylogeny inference algorithm for allele-specific SCNA data that addresses these shortcomings. MEDICC2 dispenses with the infinite sites assumption, models parallel evolution and accurately identifies clonal and subclonal WGD events. It times SCNAs relative to each other, quantifies SCNA burden in single-sample studies and infers phylogenetic trees and ancestral genomes in multi-sample or single-cell sequencing scenarios with thousands of cells.

We demonstrate MEDICC2’s ability on simulated data, real-world data of 2,778 single sample tumours from the Pan-cancer analysis of whole genomes (PCAWG), 10 bulk multi-region prostate cancer patients and two recent single-cell datasets of triple-negative breast cancer comprising several thousands of single cells.

## Introduction

Somatic copy-number alterations (SCNAs) and chromosomal instability (CIN) are hallmarks of many tumours and drive genome plasticity and intra-tumour heterogeneity (ITH) [1, 2]. SCNAs are subject to continuous evolution and selection across cancer types [3], and haplotype-resolved SCNA analyses have revealed parallel and potentially convergent evolution, including mirrored subclonal allelic imbalance (MSAI) events [4]. Besides their clinical relevance [5], SCNAs are a rich source of genetic variation that can be leveraged to reconstruct tumour evolution [6, 7]. However, for evolutionary reconstructions, SCNAs pose particular challenges, including statistical dependencies between genomic loci, overlapping of individual gain/loss events causing backmutations and physical constraints, e.g. that fully deleted genetic material cannot be regained at a later time point [6, 8, 9]. These characteristics of SCNA events necessitate an explicit evolutionary model of individual haplotype-specific copy-number changes to allow for accurate phylogenetic reconstructions.

Such an evolutionary model should also include whole-genome doubling (WGD) events [10–13], which have long been known to be linked to tumorigenesis [14–19], and which have been identified as key contributors to CIN [3, 11, 20, 21] and as potential therapeutic targets [22–24]. involves tetraploidization of genomes frequently followed by immediate loss of individual chromosomes [12, 20], thus buffering cancer genomes against the accumulation of deleterious mutations [21] and forming a substrate for further genomic diversification [3, 21]. Statistical indicators of WGD include a high average ploidy [12] in relation to the frequency of loss-of-heterozygosity (LOH) events in a cohort [25], or evidence from the clone structure of multiple samples [18]. From an evolutionary perspective, reliably detecting WGD events requires weighing a complete doubling of the genome followed by chromosomal losses against successive gains of individual chromosomes.

While several SCNA-based evolutionary inference methods have been proposed in the past [26–29], they do not model WGD events, frequently make use of the infinite sites assumption [30] and thus cannot infer parallel evolution, and do not deal with statistical dependencies between genomic loci. They are further often restricted to solving the much simpler problem of tree inference with fully sampled data, i.e. where the ancestral (internal) nodes of the tree are accessible through sequencing, an unrealistic assumption in most cases. Alternatively, many clinical studies still use hierarchical clustering based on e.g. Euclidean or Hamming distances, which are not based on evolutionary principles, to infer trees from SCNAs and interpret them as phylogenies of cancer genomes.

To address this, we have developed MEDICC2 to infer phylogenies from SCNAs based on the Minimum-Event Distance (MED) [6, 31], i.e. the minimum number of LOH events, WGD events and gains and losses of arbitrary size needed to transform one genome into another. MEDICC2 computes the MED in the presence of WGD in linear time and reconstructs phylogenetic trees, infers parallel events and ancestral genomes and times SCNA events including WGD relative to each other. We apply MEDICC2 to 2,778 tumours from the Pan-Cancer Analysis of Whole Genomes (PCAWG), where it accurately identifies WGD against a ”gold standard” set of WGD calls determined using consensus copy-number profiles from six copy-number callers [25, 32]. Using multisample prostate cancer cases we demonstrate MEDICC2’s ability to detect subclonal WGD events and to correctly place parallel evolution and MSAI events revealed by multisample phasing [3, 4]. We ultimately show how MEDICC2 infers phylogenies from allele-specific copy-number profiles for thousands of single cells without prior clustering or data aggregation.

## Results

### Inferring phylogenies from SCNAs with MEDICC2

MEDICC2 infers phylogenies and ancestral genomes from SCNAs (Figure 1A) by solving the MED problem, originally formulated by us [6] and recently studied by Zeira et al. [31], using a weighted finite-state transducer (FST) framework [33]. Briefly, the MED between a pair of copy-number profiles is defined as the minimum number of gains and losses of arbitrary length needed to transform one copy-number profile into another (see Methods). MEDICC2 thereby enforces physical constraints where gains of zero-copy segments are not permitted and zero-copy segments are ”ignored” by subsequent operations, mimicking the absence of that segment of genomic DNA (Figure 1B). This MED is thus asymmetric, and the symmetric distance between a pair of copy-number profiles is computed by minimising the MED between two copy-number profiles and their evolutionary ancestor [6] (Figure 1C). For this, the transducer implementing the MED has to be composed with its inverse, a complex operation. To avoid constructing this explicitly, we here employ a new lazy composition strategy, which only expands the FST along the path required for shortest-path computation (Methods).

**Figure 1:**
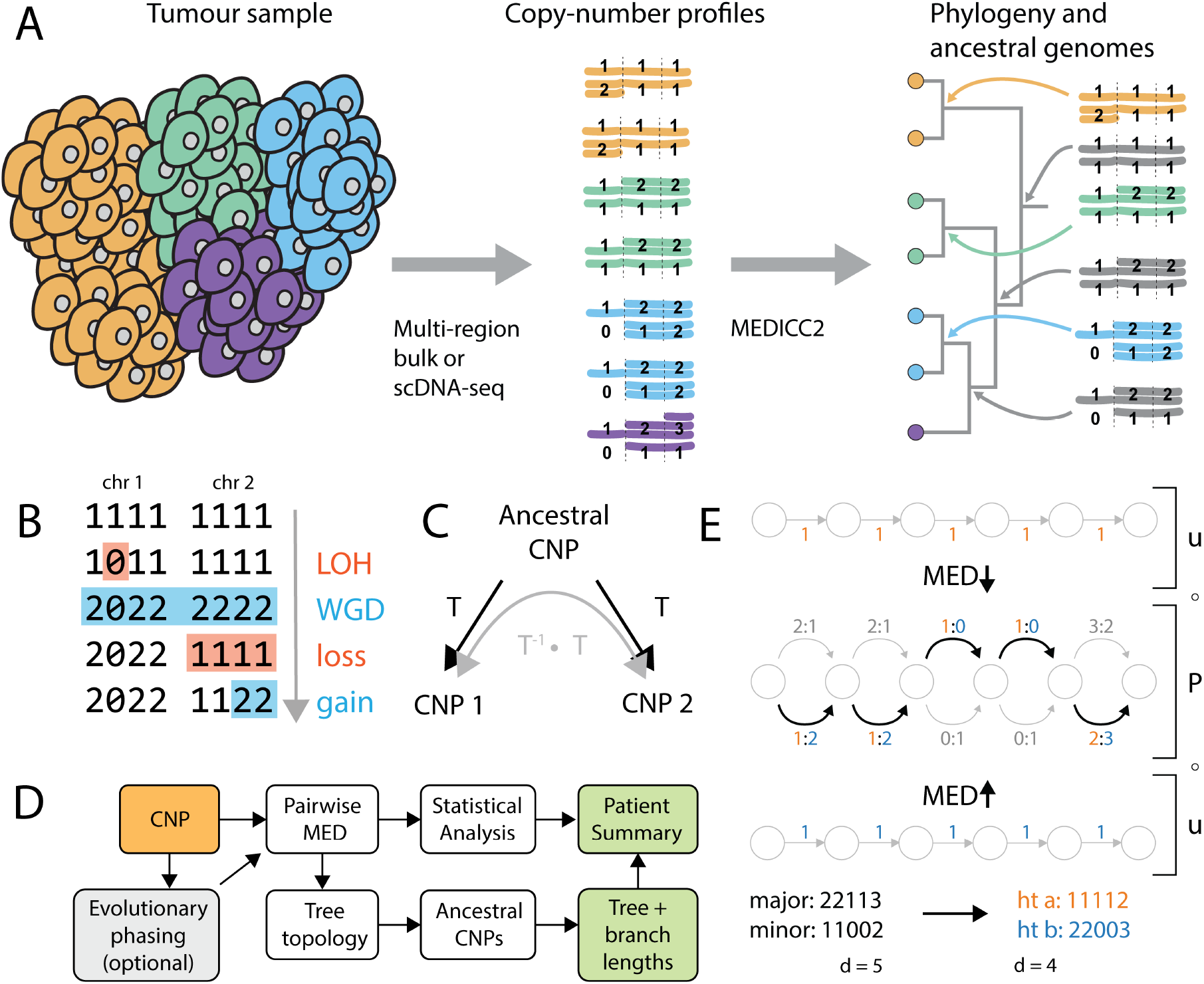
MEDICC2 algorithm. **A) MEDICC2 infers cancer phylogenies from SCNA data** from single cells or bulk sequencing using a minimum-event distance (MED) and infers the ancestral genomes. It allows for backmutations, obeys biological constraints and solves the phylogeny problem where ancestral genomes are not sampled. **B) Computing distances with WGD:** Copy-number profiles (CNPs) are represented as vectors of positive integer copy-numbers across chromosomes (here: two chromosomes with four segments each). To infer the correct MED, LOH events are considered first, as lost segments cannot be re-gained by later events. WGD events span the full CNP, whereas gain and loss events can affect an arbitrary number of segments within a chromosome. **C) Symmetric distance calculation:** The MED, as computed through the FST T, is asymmetric due to the biological constraints. The final symmetric distance is computed from a CNP to an arbitrary ancestral CNP and from the ancestral CNP back to the second CNP, thereby minimising over all possible ancestral CNPs. This is achieved by composition of the tree FST T with its inverse T^-1^. **D) Schematic overview of the MEDICC2 workflow**. Haplotype-specific CNPs are either pre-phased or undergo evolutionary phasing (see E). Pairwise MEDs are computed between all genomes, followed by tree inference and ancestral reconstruction which determines the final branch lengths of the tree. Results are reported to the user as a patient summary and plot. **E) Evolutionary phasing:** CNPs for both alleles are jointly encoded as an unweighted phasing FST P where both possible allele configurations are encoded at each position in the sequence. Evolutionary phasing then determines the optimal configuration (bold arrows) and extracts final haplotypes (orange and blue) by computing the MED between the phasing FST and two reference haplotypes (here diploid). An example of major/minor and phased configuration and its distances d to the diploid is shown at the bottom. **Abbreviations: CNP**: Copy-number profile, **FST**: Finite-state transducer, **MED**: Minimum-event distance, **LOH**: Loss of heterozygosity, **WGD**: Whole genome doubling.

To model WGD events (MED-WGD), MEDICC2 processes whole-genome copy-number profiles including both haplo-types at once, while keeping track of chromosome boundaries. Standard gain and loss events terminate when they reach the end of a chromosome, WGD events are gains applied to all non-zero segments in the genome (thereby doubling both haplotypes) irrespective of chromosome boundaries. Tetraploidization preceded/followed by rapid chromosomal loss to reach a near-triploid state [12, 20] has been described in many tumour types and is naturally contained in our model in the form of a WGD event preceded/followed by multiple losses of individual chromosomes.

Before calculating distances, copy-number profiles are typically phased (Figure 1D-E), either through the use of multi-sample reference phasing using Refphase [3], or through an internal evolutionary phasing routine (Methods), which chooses a haplotype configuration that minimises the total MED between the genome and a reference genome, typically a diploid normal (Figure 1E). MEDICC2 then infers the tree topology from pairwise MEDs between all genomes using neighbour joining [34] and calculates summary statistics as previously described [6]. Finally, ancestral copy-number profiles are reconstructed such that the total number of events along the tree is minimal, which determines the final branch lengths of the tree. The result is reported to the user as a patient summary and plot which includes the tree and inferred ancestral and terminal copy-number profiles, and change events, either globally for the whole genome or at user-defined positions of interest, e.g. oncogenes and tumor suppressors.

We first verified the technical accuracy and time complexity of the MED inference by simulating copy-number profiles with a known distance from a diploid normal under the MEDICC2 model. MEDICC2 correctly estimated the MED in linear time (Figure 2A), and the inferred MED forms a lower bound to the true number of events (minimum event criterion) with and without WGD (Figure 2B and Supplementary Figure 1A), in contrast to Euclidean distance (r^2^=0.17, Supplementary Figure 1B). The new lazy composition strategy leads to a performance increase of about one order of magnitude, enabling distance calculations for a large number of samples or single cells (Figure 2A).

**Figure 2:**
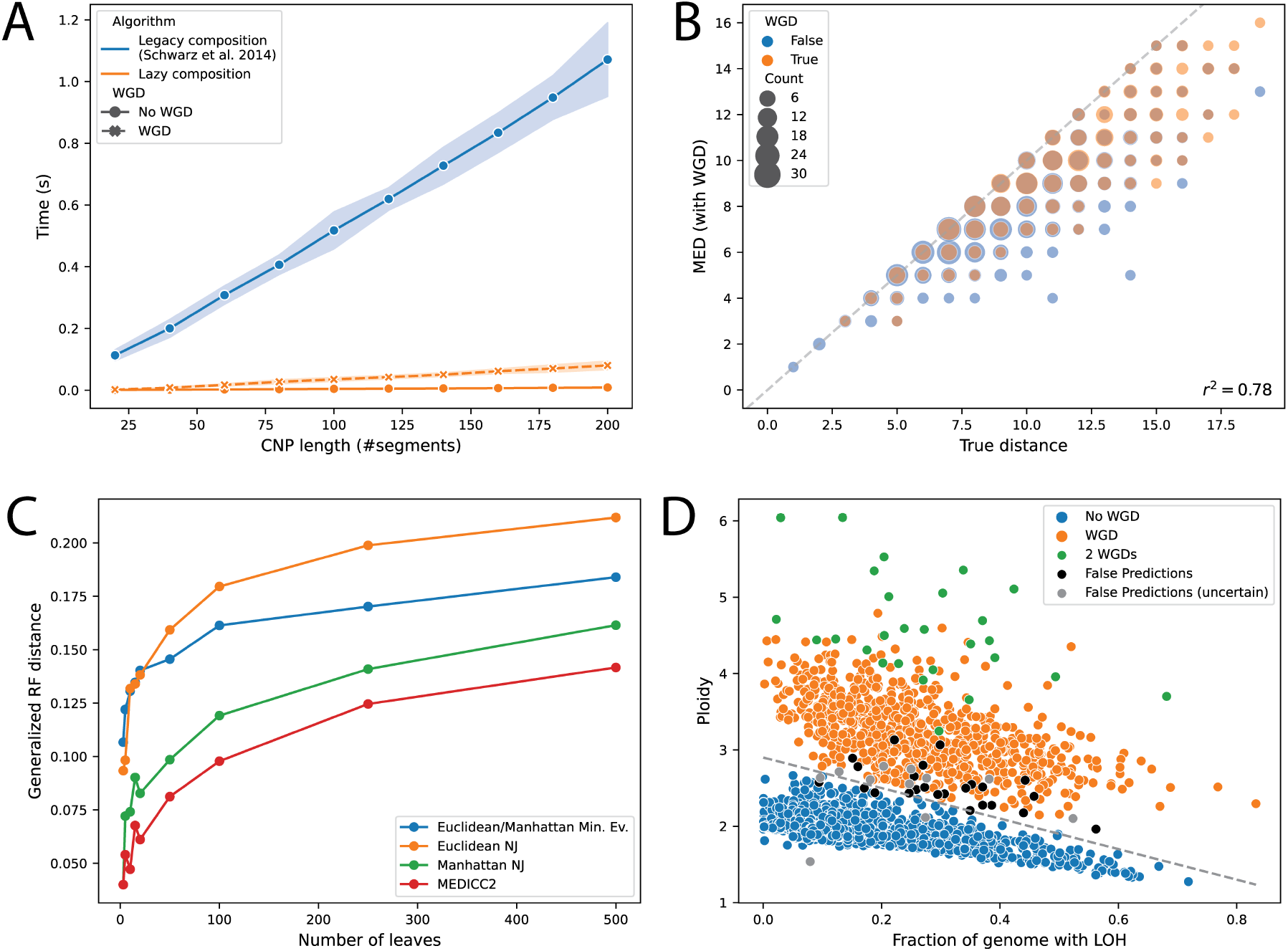
Algorithm performance: **A)** Runtime of different composition strategies for the FSTs are shown over copy-number profiles with increasing length from 20 to 200 segments. Computation time of the MEDs is linear with respect to the length of the input sequences. While MED-WGD took significantly longer to compute than the MED without WGD, the new lazy and lazy kernel composition strategies reduced runtime by a factor of between 5 and 10. **B)** Using 1000 random instances of copy-number evolution starting from a diploid genome, MEDICC2 correctly infers distances no greater than the true simulated distance if using MED-WGD (r^2^=0.87). **C)** Using an independent simulation routine we benchmark MEDICC2 against a range of other methods. The reconstructed trees were compared to the simulated trees using the Generalized Robinson-Foulds distance. As expected, the GRF distance rises with increasing tree size, while MEDICC2 outperforms all other methods for all tree sizes. **D) The MEDICC2 WGD score for 2778 cancer genomes:** Individual cancers are plotted based on their average ploidy and fraction of genome with LOH. The original separating line between WGD and non-WGD tumours was estimated by Dentro et al. as y = 2.9 - 2x. Correct ”WGD” and ”no WGD” predictions from MEDICC2 were marked in orange and blue while false predictions were marked in black and grey (latter if the PCAWG WGD status was ”uncertain”). **Abbreviations: NJ**: Neighbor-joining, **Min. Ev**.: Minimum Evolution.

We next assessed the tree reconstruction accuracy of MEDICC2 in comparison to alternative inference tools using an independent and unbiased simulation routine. Here, evolution was simulated on the level of the genome through chromosomal and segmental gains and losses but also copy-number neutral events including inversions and balanced translocations and complex events such as breakage-fusion-bridges and WGDs (Methods). From these simulated genomes with varying mutation rates copy-number profiles were generated by counting genomic segments and subjected to different tree reconstruction strategies, including Euclidean and Manhattan distances with the neighbor joining [34] and minimum evolution [35] algorithms, as well as the tailored tool MEDALT [28]. MEDICC2 outperforms other methods for all ranges of mutation rates and tree sizes, especially in the presence of WGDs (Figure 2C, Supplementary Figure 2A-B) and independent of the tree metric used (Supplementary Figure 2C).

### MEDICC2 accurately identifies WGD events in 2,778 cancers

In order to test MEDICC2’s WGD detection abilities on real data, we applied MEDICC2 to 2,778 tumours from the Pan-cancer Analysis of Whole Genomes (PCAWG) [32]. PCAWG provides high-fidelity copy-number profiles and annotations for each tumour including their WGD status, which serve us as a ”gold standard”. In PCAWG, WGD was inferred from the relationship between tumour ploidy and the percentage of the genome affected by LOH across the cohort [25]. MEDICC2 should provide a cohort-independent evolutionary measure of WGD that distinguishes between doubling of the genome, potentially followed by individual chromosomal losses, and successive individual chromosomal copy-number gains without genome doubling. To test this, we downloaded unphased consensus copy-number profiles from the PCAWG project and employed MEDICC2’s evolutionary phasing (Methods) on each copy-number profile using a diploid normal sample as a reference. We then calculated the WGD evidence score *s*_*i*_ (Methods) by comparing the MED with and without the possibility of WGD between each PCAWG tumour and a diploid normal sample. MEDICC2 prefers the presence of a WGD event over successive individual chromosomal gains and losses if the WGD evidence score is greater or equal to one (*s*_*i*_ ≥ 1). Multiple WGD events can be detected in a similar manner (Methods). For an overview of the calculated distances and the WGD evidence scores *s*_*i*_, see Supplementary Figure 3A-C.

Using this criterion, MEDICC2 predicted WGDs in 2668 out of 2778 cases (96.0%), 12 of which were predicted to have undergone two WGD events. All other 110 cases were annotated as WGD in PCAWG but these events were not called by MEDICC2 (FNR=0.04; FPR=0). Since PCAWG WGD annotations are also based on biological data with inherent noise and may contain errors, we investigated whether the 110 missed cases of WGD were marked as ’WGD uncertain’ by the PCAWG heterogeneity and evolution working group. Indeed, tumours with status ’WGD uncertain’ were significantly overrepresented amongst these tumours (*p* = 1.2 · 10^−8^, chi-square test). To increase sensitivity and in order to mitigate the effect of noisy data, we next created 100 bootstrap replicates for each sample (Methods) and calculated the WGD evidence scores for each replicate. Marking samples as WGD if at least 5% of their bootstrap runs exhibited at least one WGD event increased the detection accuracy of WGDs to 98.8% (FPR=5/2778; FNR=27/2778) (Figure 2D) while also increasing the over-representation of false predictions among tumours with status ’WGD uncertain’ (*p* = 7 · 10^−13^, chi-square test). Bootstrap sampling also identified 27 samples that underwent two successive WGDs (Figure 2D).

These results demonstrate that MEDICC2 accurately infers the presence of WGD events even in single-sample studies, without the need for additional parameter estimation or cohort-level statistics. If required, bootstrap resampling can be used to increase sensitivity and resilience against noise.

### MEDICC2 reveals subclonal WGD events and parallel evolution in prostate cancer

We next reconstructed phylogenies and inferred ancestral genomes for a multi-sample, whole-genome sequencing (WGS) cohort with 10 metastatic prostate cancer patients introduced in Gundem et al. 2015 [37]. Gundem et al. estimated cancer cell fractions of all single nucleotide variants (SNV) to perform SNV-based phylogenetic reconstructions. An illustrative example is patient ”A31” (Figure 3) which consists of one sample (”C”) from the primary tumour and four samples (”A”, ”D”, ”E”, and ”F”) from distinct metastatic sites. A31 was later included in and analysed as part of the PCAWG cohort [25] and found to demonstrate a subclonal WGD event with WGD affecting all metastatic samples (A, D, E, and F) but not the primary sample (C). In addition to faithfully recovering the original phylogeny without prior subclonal deconvolution, MEDICC2’s ancestral reconstruction correctly detected and placed the WGD event at the ancestor of the metastatic samples (WGD evidence score *s*_*A*31_ = 22, Figure 3B-C), followed by a gain on chromosome 8p and multiple chromosome wide losses. The most-recent common ancestor (MRCA) of all samples however revealed only moderate CIN with clonal LOHs on chromosomes 2, 6, 12 and 17, indicative of the substantial divergence between the primary tumor and the metastases. Finally, the ancestor of the three metastases samples A, D and F revealed a MSAI deletion on chromosome 5 different from metastasis E. Across all 10 patients four total WGD events were gathered, two of which were clonal as well as one sub-clonal and one terminal WGD (Figure 4A). We observed an overall agreement between the MEDICC2 and SNV trees, with identical topologies for 7 out of 10 patients (Supplementary Figures 4-12).

**Figure 3:**
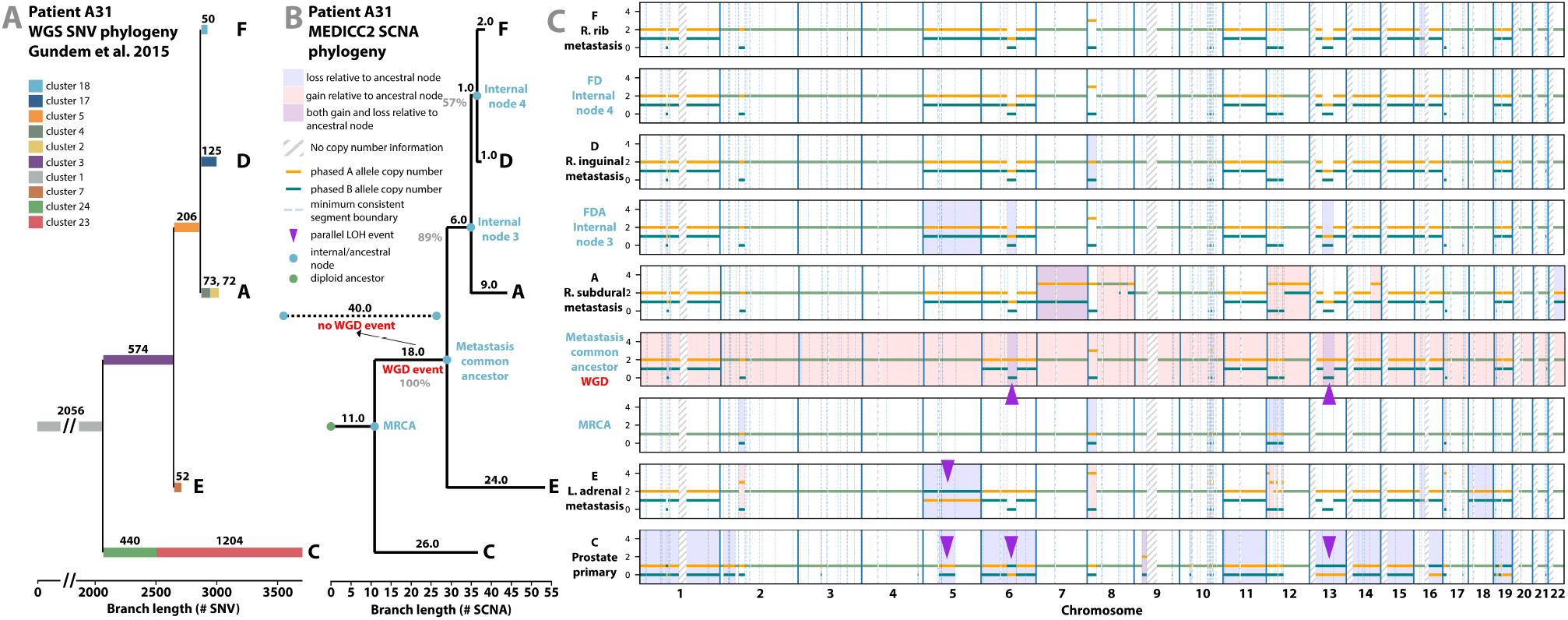
Evolutionary history of tumour subclones from patient ”A31”. **A) SNV-based phylogeny:** Reproduction of the SNV-based phylogeny as described in Gundem et al. 2015 for the multi-sample prostate cancer tumour case with one sample (”C”) from the primary tumour and four samples (”A”, ”D”, ”E”, and ”F”) from distinct metastatic sites. Original reconstruction was performed using an n-dimensional Bayesian Dirichlet process to cluster estimated cancer cell fractions of the single nucleotide variants (SNV) identified in the WGS across samples. Only the dominant clones of each sample are given. **B-C) MEDICC2 phylogeny:** Using multi-sample phased copy-number profiles, MEDICC2 detected the presence of WGD in the metastatic samples and its absence in the primary sample from A31. The MEDICC2 analysis identifies multiple MSAI events as well as parallel LOH on chromosomes 5, 6 and 13 (purple arrows). Individual events are marked in the copy-number track (C) where they occur: gains (red) and losses (blue) (see also Figure 3). The grey numbers in each branch corresponds to its bootstrap-confidence score while the events from the MEDICC2 event detection are marked in green.

**Figure 4:**
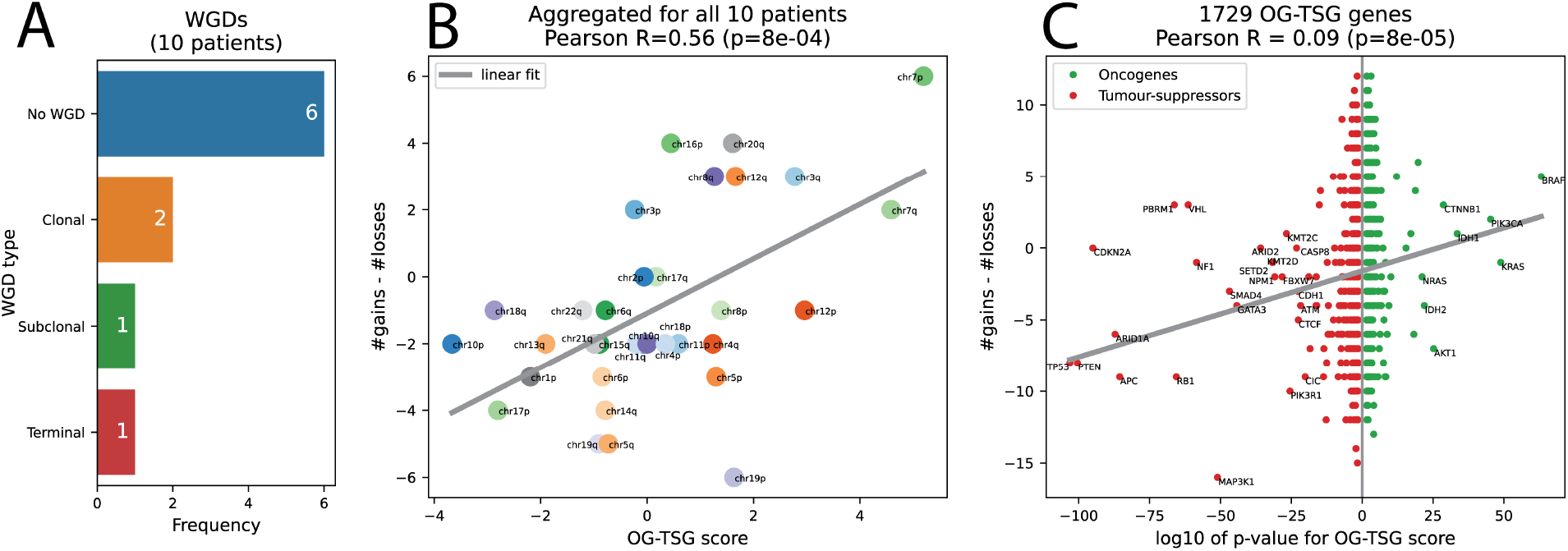
Event detection for the Gundem et al. 2015 cohort. A) WGD detection: In the 10 patients a total of 10 WGDs were detected, two of which were clonal, one sub-clonal and one terminal. **B)** Using the MEDICC2 event detection routine we detected the number of times a whole chromosome arm was either gained or lost completely in a single branch. The gains and losses were aggregated over all patients and samples into a -single score. This score was compared against the oncogene - tumor suppressor gene (OG-TSG) score derived by Davoli et al. [36] A clear correlation between the gains/losses and the OG-TSG score (which is not based on copy-numbers) is visible. **C)** The analysis was repeated on the basis of all 1729 individual genes present in the Davoli et al. dataset. On the x-axis we plotted the base-10 logarithm of the genes’ p-value and flipped the sign for the oncogenes to create a single, continuous x-axis for both genesets. A small correlation is visible which becomes more pronounced when only considering the top 100 genes. Genes with *p* < 10^−20^were marked with their name.

We next compared relative branch lengths within the SNV-based tree of A31 and the SCNA-based MEDICC2 tree. Branch lengths of both trees correspond to the number of SNVs present in each subclonal cluster and to the number of SCNA events larger than 1 Mb, respectively. To facilitate comparison between branch lengths in the two trees we computed relative event distances by normalising branch lengths by the maximal root to leaf distance for each tree. The relative event distances from the root of each tree to the terminal nodes representing samples A, F and D were conserved in the two phylogenetic reconstructions. However, the SNV-based and SCNA-based trees demonstrated distinctly different relative branch lengths from the diploid normal root leading to the MRCA. This ”trunk” was found to be shorter in the SCNA-based MEDICC2 tree (11/53 SCNAs) when compared to the SNV-based tree (2056/2682 SNVs). This suggests that there have been relatively few founder SCNAs compared to a large number of founder SNVs, potentially due to a larger number of SNVs present in the tissue before malignant transformation and a later onset of CIN likely as a result of one or more of these SNVs. This finding was replicated in 9/10 patients of the full cohort (Supplementary Figures 4-12). In addition, in A31 the branch terminating at the dominant clone of metastatic sample E was relatively long in the SCNA-based tree compared to in the SNV tree (24/53 vs 52/2682, Figure 3B). This relatively large minimum event distance is due to multiple SCNA events on chromosome 12 present only in sample E suggesting a complex set of SCNAs potentially occurring together.

Recently, we applied the multi-sample reference phasing algorithm that maintains consistent phased haplotypes across samples from a single patient’s disease to reveal additional SCNA heterogeneity across human cancers [3, 4]. This additional heterogeneity results from the detection of MSAI as well as SCNA-mediated parallel evolution where the same SCNA event (e.g. an LOH event) occurs independently affecting distinct haplotypes within an individual patient’s disease [3, 4, 38, 39]. Since MEDICC2 models both haplotypes individually and does not employ the infinite sites assumption, it can infer both MSAI-mediated homoplasy and homoplasy affecting the same allele by assigning these parallel events to separate branches of the tree. Multi-sample reference phasing analysis of the samples from A31 identified multiple MSAI events as well as parallel LOH on chromosomes 6 and 13 (Figure 3C). MEDICC2 assigned the independent origins of these parallel events to the branch corresponding to the emergence of the dominant clone in the primary sample and to the branch corresponding to emergence of the common ancestor of all the metastatic samples (Figure 3B-C). MEDICC2’s ability to correctly identify and locate these parallel evolutionary events revealed by multi-sample phasing provides additional evidence for a diverging evolutionary trajectory between primary and metastatic samples, absent from its original analysis [37].

We were further interested in whether the inferred tree topologies and SCNAs can be used to detect preferentially gained and lost regions, potential indicators of positive selection [3]. To this end we used the oncogene (OG) and tumor suppressor gene (TSG) scores derived by Davoli et al. [36] for individual genes as well as on the level of chromosome-arms. MEDICC2’s event detection algorithm (Methods) allows calculating the net number of gains and losses along the phylogenetic tree in regions of interest and counts events only once at the node in the tree where they occur. Across the 10 patients in this cohort, a clear correlation is visible (Pearson R=0.48, p=0.0053) between the MEDICC2 event score and the OG-TSG score on the level of chromosome arms reported by Davoli et al. [36] (Figure 4B, Supplementary Figure 3B). Additionally, on the level of all 1,729 individual genes from Davoli et al., we find a correlation of R=0.09 (p<0.0001) (Figure 4C) which rises to R=0.27 when considering only the top 100 genes. Despite the low sample size of 10 patients, the results show the ability of MEDICC2 to infer regions of interest by detecting distinct gain and loss events in the individual copy-number trees.

### MEDICC2 infers SCNA phylogenies from single-cells

Recent advances in single-cell technology have enabled the collection of copy-number profiles of thousands of cells. While large single-cell experiments constitute a major opportunity to study tumour evolution with higher precision and on a larger scale, they also bring unique challenges. The lower coverage of single-cell studies lead to a lower signal-to-noise ratio than conventional methods and therefore to less reliable and more noisy copy-number profiles. The large number of copy-number profiles representing cells increases the computational burden, in particular for pairwise distance calculations and ancestral reconstruction. Due to the new fast composition algorithm (Figure 2A) and an efficient parallelisation strategy (see Methods and [40]) MEDICC2 processes thousands of cells efficiently. Here, we apply MEDICC2 to a previously published single-cell study of triple-negative breast cancer by Minussi et al. [41] looking at the two patients highlighted in the paper, TN1 and TN2, with 1,100 and 1,024 cells, respectively. In the original study, the authors defined ”superclones” and ”subclones” by two separate clustering methods in the two-dimensional UMAP space created from pairwise Manhattan-distances. Consensus copy-number profiles were created from these clusters and a minimum evolution tree was created from the Manhattan distances between these consensus profiles. This indirect way of determining the phylogeny of these cells involved a number of data abstractions that involved manual selection of hyperparameters (e.g. for the clustering algorithms).

We instead derived allele-specific copy-numbers from the original raw data (Methods) and ran MEDICC2 directly on the allele-specific copy-number profiles to reconstruct phylogenies for all cells without intermediate clustering steps or consensus profiles (Figure 5 and Supplementary Figure 13A).

**Figure 5:**
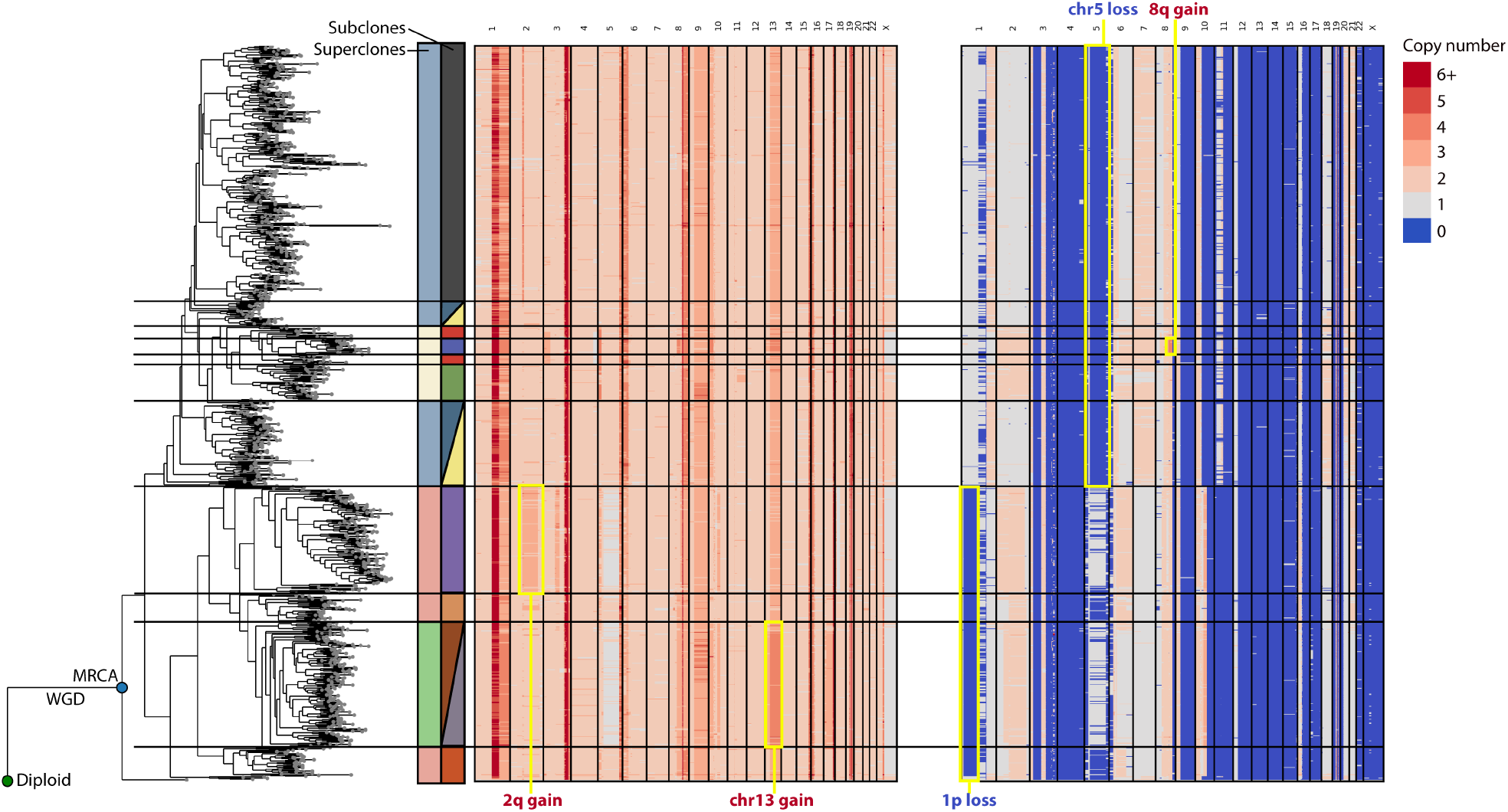
Inferred phylogeny for single-cell data with 1,024 cells. Inferred phylogeny and allele-specific copy-number profiles for patient TN2 from Minussi et al. 2021. The diploid and most recent common ancestor to all cells are marked with green and blue circles, respectively. We manually selected clades from the phylogeny to match the superclones and subclones of the original publication. These are marked next to the tree in the colors of the original publication and with horizontal lines. The structure of the tree corresponds very clearly with distinct features of the copy-number profiles and match the clonal structure derived in the original publication. Selected synapomorphies of the clone structure are highlighted with a yellow border and annotated on the figure.

Next, we mapped superclones and subclones from the original publication to the MEDICC2 tree and found a high degree of concordance between the clonal architecture revealed by MEDICC2 and the original results [41], in contrast to a simple tree based on Manhattan distance between all cells (Supplementary Figure 13B-C). For TN2, MEDICC2 recreates all superclones and most subclones from the original publication, while for TN1 it consolidates two superclones into one, but otherwise detects them as in the original publication. In addition, MEDICC2 correctly detected truncal WGDs in both patients as described [41].

While the original study reports truncal branch lengths similar to the maximal MRCA to leaf distance, suggesting that roughly half of the SCNA events happened before emergence of the MRC, we find truncal branch lengths substantially shorter in the MEDICC2 phylogenies (42/164 for patient TN1 and 71/238 for patient TN2). These findings are in concordance with our results for the metastatic prostate cancer patients described above and provides further evidence for substantial clonal diversification after emergence of the MRCA.

Our analyses demonstrate that MEDICC2 infers tree topologies that provide substantial biological insight, while previous approaches using general measures such as the Manhattan distance were not able to recover the clonal architecture of the tumour (Supplementary Figure 13B-C). In contrast to clustering of consensus profiles, MEDICC2 retains single-cell information when inferring tree topologies and ancestral genomes. To the best of our knowledge, MEDICC2 is the only available algorithm that can reliably create accurate phylogenies from thousands of single-cells.

## Discussion

We here develop and apply a computational approach for reconstructing the evolutionary history of cancer from haplotype-specific SCNA profiles. MEDICC2 employs an explicit evolutionary model of copy-number change, computes the MED [6] in linear time at a fraction of its original runtime. It detects individual SCNA events, including clonal and subclonal WGD, and provides statistical robustness assessment of the inferred trees. MEDICC2 is applicable to any allele-specific or total copy-number estimation algorithm from any sequencing modality or SNP array. In contrast to breakpoint-based approaches, MEDICC2 models actual genomic events that change copy-number, incorporating biological constraints such as in LOH regions, where the lost allele cannot be regained.

MEDICC2 does not model the underlying biological processes, such as whole chromosome missegregation [2], chromothripsis [42], chromoplexy [43], breakage fusion bridge cycles [44] and others [45]. In the future such events might be possible to incorporate depending on the additional computational complexity required. Nonetheless, our simulations show that MEDICC2 accurately infers phylogenies in the presence of these complex events, outcompeting alternative methods. In these simulations, the recently developed MEDALT that also incorporates MED, performed poorly relative to the other methods considered (Supplementary Figure 2B). One possible explanation might be that, like many other tools, MEDALT uses a minimum spanning tree to infer the tree topology, thereby implicitly assuming that the set of sampled genomes also contains the ancestral genomes from the tumour’s past.

Our recent work [3, 4] and that of others [38] has highlighted the importance of using multi-sample phasing to reveal additional SCNA heterogeneity taking the form of MSAI or parallel evolution of similar SCNAs from distinct haplotypes [3, 4]. Since MEDICC2 does not employ the infinite sites assumption, it can reveal homoplasy on alternating haplotypes (MSAI) as well as on the same haplotype where an independent origin of two events leads to a more parsimonious phylogeny than a shared ancestry.

WGD in cancer can result from endoreduplication [46], mitotic dysfunction [47] or cytokinesis failure [48]. Nearly all of these proposed mechanisms suggest a diploid to tetraploid transition (tetraploidization) [49], frequently followed by return to a near-triploid state through subsequent chromosome losses [20], possibly with preceding LOH events [50]. MEDICC2 can replicate this behavior naturally through the combination of LOH events, a WGD event and multiple independent chromosome-wide losses. As every chromosome-loss is counted as a separate event, the MED might over-estimate the true number of events. However, our model seems to fit real-world copy-number profiles extremely well as verified on 2,778 WGS tumours from the PCAWG cohort.

In the Gundem et al. cohort [37] MEDICC2 provided further support for the divergence of the primary and metastatic samples through the detection of a subclonal WGD event and parallel evolution in A31. Interestingly, other SCNA-mediated parallel evolution events were identified at the *AR* gene locus in the original analysis of this tumour through the orthogonal method of exact structural variant breakpoint identification enabled by WGS [37]. Analysis of the net gains and losses of chromosome arms and individual genes along the inferred trees for all patients showed a clear correlation with the OG-TSG score [36]. This demonstrates the ability of MEDICC2 to find genomic events potentially under positive selection for clinical interpretation of tumour evolution. In the future, analysis of a larger cohort could yield further insights into preferentially gained and lost regions of the genome in a cancer-type specific way.

In contrast to SNV data, assessing within-sample subclonality from copy-number profiles from bulk sequencing is notoriously difficult and bulk copy-number profiles thus typically only reflect the dominant clone in that sample. Since MEDICC2 is agnostic of the sequencing modality used to generate its input, future advances that increase the resolution of copy-number profiles will be usable by MEDICC2 without modification. Here, single-cell sequencing [51] for total [52] and allele-specific copy-number profiles [53] can help increase resolution. As we have shown, MEDICC2 can handle thousands of cells and thereby infer inter- and intra-region evolution. It clearly outperforms pairwise Manhattan distances from the original study, creating a tree topology that matches previously identified super- and subclones with high accuracy directly from single cells without additional parameter fitting or the creation of consensus profiles.

Like most computational tools, MEDICC2’s performance is limited by the quality of the input data. Over-segmentation, a common problem of copy-number inference, can have large effects on the MED and this issue is elevated in single-cell experiments which are prone to low signal-to-noise ratios. In the future, co-inference of copy-number and tree topology will help resolve ambiguities due to noise in the data [54].

In summary, systematically determining the number and order of WGD, arm-level SCNAs, and focal events that have occurred in the evolutionary history of a tumour has not yet been performed on a large scale and has previously been the preserve of theoretical mathematical modelling [55, 56]. MEDICC2 enables the reconstruction and timing of the individual SCNA events present in the evolutionary history of a tumour that may overlap and build upon one another. This will allow much more detailed dissection of WGD, aneuploidy and CIN across the genome utilising single sample, multi-sample and single-cell approaches, than the measures of the proportion of the genome affected by SCNA that much of the field has previously relied on.

## Methods

### The MEDICC2 model

To solve the MED problem we employ a finite-state transducer (FST) framework as previously described [6], following the notation of Mohri [57]. Copy-number profiles are represented as vectors of positive integer copy-numbers (*k*)_1..*n*_, with 0 ≤ k ≤ 8, where each integer copy-number represents a genomic segment *i*. Chromosome boundaries are marked by a chromosome separator character ’X’ and both haplotypes are concatenated and separated by ’X’. We represent these allele-specific copy-number profiles as unweighted finite-state acceptors (FSA) *A* = (Σ, *Q, E, i, F*) and evolutionary events as weighted FSTs *T* = (Σ, *Q, E, i, F, λ, ρ*) with (input and output) copy-number alphabet Σ = {0, .., 8, *X*} (per allele), a finite set of states *Q*, a finite set of transitions *E*, an initial state i ∈ *Q*, a set of final states F ⊆ *Q*, an initial weight *λ* and a final weight *ρ*. Transitions between states are equipped with an input symbol *l*_*i*_ ∈ Σ(input copy-number) and an output symbol *l*_*o*_ ∈ Σ (output copy-number) and a weight *w*. All weights *λ, ρ, w* are taken from the positive integers including zero and calculations are carried out over the tropical semiring, i.e. weights are summed along the path of a FST and the final weight between a pair of sequences is the minimum over all possible paths

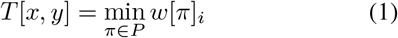

(see [33]), where *P* is the set of all possible paths transforming *x* to *y*.

FSTs and FSAs can be subjected to a variety of operations, of which *composition* (”◦”) is of particular importance. During composition a new FST is constructed in which the set of states is the cartesian product of the set of states of the two input FSTs. The composition *S* of two FSTs *T*_1_ and *T*_2_ then assigns a weight to any pair of input and output sequences by chaining their transduction

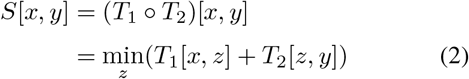

via intermediary sequence *z* [57]. Composition is also used to effectively compute the score or total weight *T* [*x, y*] that a FST *T* assigns to a pair of sequences *x* and *y* (Eq. 1) by representing *x* and *y* as two unweighted acceptors and running a single-source shortest distance algorithm (SD) over the composition x ◦ T ◦ y [57]

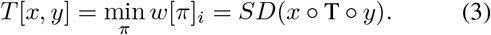

Composition enables us to combine multiple evolutionary event FSTs into a final FST in which the individual events are carried out successively in order of composition, and to transform the asymmetric MED into a symmetric MED for calculation of the pairwise distance matrix [6].

### Calculating the minimum-event distance

It has been shown previously that the standard MED can be solved by considering losses separately before any gains [31]. Indeed, only loss-of-heterozygosity (LOH) events, i.e. losses which reduce haplotype-specific copy-numbers to zero, must be considered first, as subsequent gain and loss events must ignore the positions with copy-number zero. The MED however is oblivious to the ordering of any subsequent gains and losses. When including WGD events (MED-WGD), LOH events must again be dealt with before any other event. In addition, WGD events must come before any segmental losses and gains, for example to allow for the deletion of segments previously gained during a WGD event (Figure 1B). The inclusion of WGD events further introduces non-determinism into the problem as locally WGD events cannot be distinguished from segmental gains before taking the full sequence into consideration.

We thus define four one-step FSTs which model one of four different evolutionary events considered: (i) LOH events (*T* ^1^_LOH_), (ii) segmental (+1) gains (*T* ^1^_*G*_), (iii) segmental (−1) losses (*T* ^1^_*L*_) without LOH and (iv) WGD (+1 for all non-zero segments) events (*T* ^1^_WGD_). LOH events, gains and losses must terminate when they reach separator character ’X’. WGD events do not terminate at ’X’ and leave it unchanged. In the one-step FSTs, each sequence position can only be affected by a single event. For example, the one-step FST for segmental gains *T* ^1^_*G*_ only allows copy-number changes of arbitrary length from 1 to 2, 2 to 3, and so on, but not, for example, from 1 to 3. To span the full range of possible events, the one-step FSTs are each composed *n* times with themselves and the maximum copy-number dictates the number of compositions necessary: *n* = |Σ| − 1 for LOH events and *n* = |Σ| - 2 for segmental losses and gains and WGD events [6]. The resulting event FSTs *T*_LOH_, *T*_*G*_, *T*_*L*_, and *T*_WGD_ are then chained (composed) into the asymmetric MED-WGD FST)

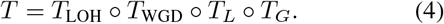

The final MED-WGD between copy-number profiles *x* and *y* is then computed following Eq. 3 as

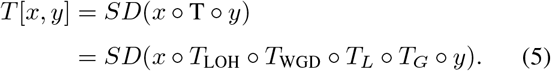

Analogously, the simple MED is built via composition as in

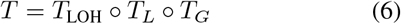

and distance calculation is carried out as in Eq. 5.

As noted previously the MED and MED-WGD are asymmetric. To compute symmetric distances *S*[*x, y*] between pairs of copy-number profiles connected in a phylogenetic tree we compute the score between *x* and *y* via its common ancestor using the kernel composition of *T* with its inverse *T* ^−1^ [58, 59] (Figure 1C):

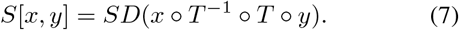

As the number of states in a composed FST is the product of the states of the input FSTs, explicit computation of the composition in Eq. 7 is computationally expensive. We therefore employ a new computation strategy based on lazy (on-demand) composition followed by shortest path computation using a shortest-first queue [60]. Lazy composition prevents full expansion of the composed FST before determining the shortest path and instead expands the FST only along the path visited [60].

### MED speed and accuracy evaluation

To assess the performance of the new lazy composition strategy and the accuracy of the MED calculation we simulated copy-number profiles following the MEDICC2 evolutionary model. A random number of evolutionary events was generated using a poisson process with rate parameter *µ* = 10 (reconstruction accuracy test) and *µ* = 20 (speed test). In the reconstruction accuracy test each event had a 5% probability to be a WGD event, and a 47.5% probability of being a gain or loss respectively. In the speed test, to prevent too many deletions, the gain probability was set to 80%. The start of an event of gain or loss events in the sequence The start of an event was selected uniformly at random from the set of remaining available positions (positions with copy-number ≠ 0) and event lengths were drawn from a geometric distribution with success probability parameter *p* = 0.2. Events were applied to the sequence obeying biological constraints, i.e. no gain of segments with copy-number zero and forced ending of events at chromosome boundaries, the latter with the exception of WGD. For the reconstruction accuracy test, sequences were fixed at length 50 (five chromosomes of length 10 each). For the speed test, sequence lengths were varied from 20 to 200 segments (Figure 2A).

### Linear time evolutionary phasing

Traditionally, allele-specific SCNAs are reported in major and minor copy-number, as the relative phasing of copy-number segments to each other is unknown. We introduced the multi-sample reference phasing implementation *refphase* [3, 61] to leverage relative phasing information in a multi-sample sequencing scenario and used it to identify MSAI events across human cancers [3, 4, 39]. In situations where multi-sample reference phasing is not feasible, e.g. in single-sample scenarios, we developed evolutionary phasing [6], where the assignment of major and minor copy-numbers to parental haplotypes is chosen to minimize the sum of MEDs over both parental haplotypes (minimum evolution criterion). In its original form, evolutionary phasing was achieved through the use of a weighted context-free grammar in concert with our original MED [6], a computationally costly solution. To enable phasing for a large number of segments and genomes, we here provide a novel phasing strategy which solves the evolutionary phasing problem exactly, but at a fraction of the original runtime, by staying within the realm of regular grammars and FSAs.

To do so we first encode copy-number profiles for both alleles jointly as an unweighted phasing FST *P* as follows: the FST follows a linear structure with a number of states equal to the number of segments +1. Two transitions occur between each neighboring pair of states and the two transitions have as input symbols major copy-numbers and as output symbols minor copy-numbers and vice versa (Figure 1E). Due to these mirrored input and output symbols every valid path through the phasing FST *P*thus determines an assignment of copy-number alleles to haplotype 1 and haplotype 2. The set of of all 2^*n*^possible paths for a sequence of length *n* through this FST corresponds to the set of possible phasing choices. To choose the most parsimonious haplotype assignment, this phasing FST is then composed from the left and from the right side with a composed FSA *u* = (*d* ◦ *T*) ↓of the diploid FSA d(encoding all-1s) with the MED-WGD FST *T*, projected to its output (↓). Shortest-path (SP) computation over this composite yields the optimal phase with a total score equal to the sum of MEDs over both parental haplotypes. Separate haplotypes *h*_*a*_and *h*_*b*_ can be extracted by projection to input and output followed by weight removal:

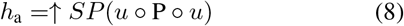

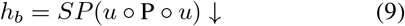

### Simulating genome evolution

To evaluate the performance of MEDICC’s tree reconstruction algorithm first a tree topology for a given number of leaves was created by randomly joining sample labels and rooting the tree at the diploid. The branch lengths and therefore the number of events per branch were determined using a Poisson distribution with *λ* = Δt · S · *µ*, where Δ*t* was set to 1, *S* represents the length of the genome (440 segments, see below) and *µ* a variable rate parameter. To avoid biases, somatic evolution was modelled on the level of the genome, not the copy-number level, along the tree starting at the diploid. We chose 2 × 22 chromosomes (two sets of haplotypes) with 10 segments of uniform size each which resemble the makeup of many actual bulk copy-number profiles. At every branch, the genome was mutated with a number of genomic events based on the corresponding branch length. These events encompass gains and losses of whole chromosomes, focal losses, insertions, breakage-fusion-bridges (BFB), whole genome doublings (WGD) as well as copy-number neutral events such as balanced and unbalanced translocations and inversions. For example, if a segment from chromosome 1 is moved to chromosome 2 through an unbalanced translocation and chromosome 2 is subsequently gained, the segment of chromosome 1 is also gained. By choosing this approach we ensure that the simulation is not biased towards the approach of MEDICC2 (which rather models the evolution of copy-number profiles and not individual segments) and mirrors actual tumor evolution. In the absence of actual event probabilities we kept all events to be equally likely with the exception of BFBs and WGDs. The probability of BFBs was set to ¼ of the other events and for the WGD we choose four different probabilities: 0.000125 for the simulation of large trees reminiscent of single-cell experiments (Figure 2C), and three levels for the simulation of medium sized-trees (0 for ”No WGD”, 0.0125 for ”Low WGD” and 0.065 for ”High WGD”, (Supplementary Figure 2B).

For the large tree scenario we simulated 25 trees each for all combinations of the mutation rate *µ* ∈ [0.01, 0.025, 0.05]and the number of leaves N ∈ [5, 10, 15, 20, 50, 100, 250, 500]. For the medium tree scenario we simulated 25 trees each for all combinations of the mutation rate *µ* ∈ [0.01, 0.025, 0.05, 0.075, 0.1]and the number of leaves N ∈ [5, 10, 15, 20]and the three levels of WGD as described above.

Reconstructed trees were evaluated using the Generalized Robinson-Foulds (GRF) distance as implemented in the R package TreeDist [62]. The GRF is based on the widely used Robinson-Foulds distance which measures the number of splits that occur in both trees. The GRF improves this metric by taking the similarity of splits that are not perfect matches into account. We furthermore used the regular Robinson-Foulds distance (as implemented in the R package ape [63]) and the Quartet distance (as implemented in the R package *Quartet*) to prevent any potential biases from the tree metric used (Supplementary Figure 2C).

We compared MEDICC2 to a range of widely used methods which encompassed Euclidean- and Hamming-distance based trees created both through neighbor joining and minimum evolution. For neighbor-joining we used the implementation of MEDICC2 and for the minimum evolution tree we used the function *fastme*.*bal* from the R package *ape* [63]. As a representative of algorithms that create minimum spanning trees (MST), we also compared against MEDALT [28] (Supplementary Figure 2A).

### WGD detection

To facilitate WGD detection, we calculate the MED with (MED_WGD_) and without the possibility of WGD (MED_noWGD_) between each PCAWG tumour and a diploid normal sample and computed the WGD evidence score *s*_*i*_ as

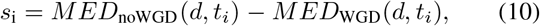

where *t*_*i*_ represents a PCAWG tumour profile and *d* represents a standard diploid normal sample. Because ME*D*_noWGD_(*x, y*) *> MED*_WGD_(*x, y*) for any valid set of copy-number profiles *x* and *y*, the score *s*_*i*_ is always positive (*s*_*i*_ ≥ 0) and a score of *s*_i_ ≥ 1indicates a preference for a WGD event to have occurred.

By replacing the multi-step WGD transducer *T*_WGD_ in (Eq. 4) with n-step WGD transducers for variable n, we can test for multiple WGD events. For example, the scores ME*D*_1*W GD*_(*x, y*) *> MED*_2*W GDs*_(*x, y*) = *MED*_3*W GDs*_(*x, y*), indicate two WGDs to have taken place.

In order to increase the robustness of our predictions, we repeated the analysis with 100 bootstrap runs (see below). Samples that exhibited WGDs (or multiple WGDs) in at least 5% of the bootstrap runs were classified as WGDs (or multiple WGDs, respectively).

### Event detection and correlation with OG-TSG score

For comparisons between events detected in MEDICC2 and the OG-TSG score we downloaded 1729 gene annotations from Davoli et al. [36] and the aggregated, chromosome-arm wise OG-TSG scores that measures the occurrences of OGs and TSGs on a given arm. To extract events we leverage the ancestral reconstruction routine in MEDICC2. Trees are then traversed in postorder, relative copy-number changes are determined for all segments and events are counted in the branch where the change occurs, thereby taking parallel evolution into account while preventing counting the same event multiple times in multiple samples from the same patient. Change events were then overlapped with regions of interest, i.e. the positions of OGs and TSGs as well as the chromosome arms. An event is detected if there is at least 90% overlap between the event and the region of interest. Gains and losses are summed across all branches and patients to arrive at the final ”#gains - #losses” score for each gene / chromosome arm. The event detection routine is available to MEDICC2 users by providing BED files with regions of interest and MEDICC2 can calculate the number and exact location of gains/losses of these regions along the evolutionary trajectory.

### Resampling for robustness estimation

The bootstrap [64, 65] is a classical approach in phylogenetics to assess the robustness of an inferred tree to perturbations of the data. During bootstrapping of a multiple sequence alignment columns are drawn from the original data with replacement and a large number of resampled datasets (typically 100-1000) are created. The tree reconstruction method of choice is then employed on all bootstrap datasets and the relative frequency with which a branch (or taxon split) of the original tree appears in the set of bootstrapped trees forms a support value for this branch. A necessary requirement for this approach is the independence of sites in the alignment. Since this assumption does not hold for copy-number profiles, we use the following alternative resampling strategies for copy-number profiles in MEDICC2:

#### Chromosome-wise bootstrap

Here, whole chromosomes are drawn with replacement from the original chromosomes to create a bootstrap sample. As losses and gains end at chromosome boundaries and as WGD events are ignorant to the order and number of chromosomes, this approach does not introduce false events while still providing a sufficiently large sample space, albeit at the cost of a coarse-grained resolution. Therefore not all bootstrap samples will be equally representative of the underlying data.

#### Segment-wise natural jackknife

Here, N segments are drawn with replacement from the original N segments, discarding all duplicates. On average this is equivalent to discarding 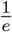 randomly selected segments [66]. The jackknife approaches the bootstrap distribution and due to the lower number of resulting segments has a speed-advantage over the chromosome-wise bootstrap, however, the jackknife generally generates less accurate representations of the original data than the bootstrap. Branch support values are indicated by their percentage value on the respective branches (see Figure 3A).

### Parallelization strategy

Single-cell experiments with thousands of cells demand high performing methods as the pairwise distance calculations scale with *O*(*N*^2^) and are therefore exceptionally computationally expensive. In addition to the performance improvements when calculating the MED, we implemented a parallelization routine to make MEDICC2 applicable to 100s to thousands of cells. To this end we utilized a recently proposed parallelization strategy [40] to split the *N* × *N* pairwise distance calculations into smaller chunks that can be run in parallel. In the method used, the N samples are split into *p*^2^ + *p* groups of size p (where p is the smallest prime such that *p*^2^ ≥ *N*) and the pairwise distances within the individual groups are calculated such that a given pair is never calculated twice. This allows for a theoretical speed-up by the factor *p*^2^ + *p* which for all practical concerns is only limited by the number of available cores [40].

### Single-cell data processing and analysis

Segmented logged ratios of read counts within genomic bins and total copy-number profiles of single-cell triple-negative breast cancer data were obtained from ref [41]. Allele-counts at 1000G SNP positions were obtained for each single cell using alleleCounter (v.4.0.0) as described in ref [41].

#### Fitting to integers

The logged ratios Logged ratios were centred to zero by subtracting the mean to obtain the logR. logR values were fitted to integers by identifying the offset *ψ*that minimises the sum of distances across segments of the n_tot_=logR-*ψ* to their values rounded to the closest integers round(n_tot_), weighted by the lengths of the segments w: argmin _*ψ*_ *w*_*i*_ × (*n*_*tot,i*_ — *round*(*n*_*tot,i*_))^2^|*n*_*tot,i*_ = *log*(*R*_*i*_) − *ψ*. In the original publication, the logged ratios were fitted to integers by using the same FACS ploidy value as the offset for all cells. Since individual cells can harbour private CNAs, their ploidy can indeed vary around their average FACS ploidy. Therefore, we derived the average number of copies along the genome calculated from the published total copy-number profiles (∼ initial value of *ψ*according to FACS ploidy) and performed a search within {0.85*ψ*, 1.15*ψ*} by steps of 0.01 to further optimise the offset to minimise the distance to integers within each individual cell.

#### Getting phased parental-allele-specific copy-number profiles. Identify heterozygous SNPs

Across all cells from the same patient, allele counts were summed to get a pseudo-bulk profile. FACS sorting based on ploidy enriches for tumour cells, but still 10-15% of cells were normal contaminants [41], thus even in LOH regions, heterozygous SNPs can be identified. As described in ref [41], heterozygous SNPs with allele counts for genotype A and B, c_A_ and c_B_, were defined as those with P(Bin(c_A_ + c_B_, 0.99) ≤ c_A_) < 0.01 and P(Bin(c_A_ + c_B_, 0.99) ≤ c_B_) < 0.01. At each heterozygous SNP position, the genotype with the highest read count in the pseudo bulk was assigned to the major allele.

#### Fitting within cells

After phasing all heterozygous SNPs, for each segment, the maximum likelihood estimate of the BAF b_mle_ is derived as follows: from each b belong to the possible values between 0 and 1 by steps of 0.001, b_mle_ is the value of the BAF b that maximizes the likelihood of a Binomial distribution with probability b, number of successes is the total number of reads bearing the genotypes assigned to the major allele, and the number trials is the total number of reads.

#### Fitting across cells

To account for the noise in n_tot_ and BAF, copy-number states of each segment are assigned by fitting these data to integers across cells. Each cell’s segment is assigned to allele-specific copy-number states as follows: first, it is assigned to its closest integer allele-specific copy-number state, i.e. {round(n_tot_*BAF), round(n_tot_)- round(n_tot_*BAF)}; second, at each populated allele-specific copy-number state across cells, the noise parameter for a Gaussian distribution is estimated from the non-rounded integers, with the mean being the total integer corresponding to the integer state, and the parameters for a Beta distribution are estimated from the segments’ BAF values, keeping the mean of the Beta as the BAF of the corresponding integer state; then, each cell’s segment is re-assigned to the allele-specific copy-number states that minimise the sum of its LogR and BAF likelihoods normalized across states; the weight given to the likelihood from the LogR can be modulated to best assign states from diploid cells ((1.9<ploidy<2.1) to {1,1} across segments (here, 50% more weight was given to the likelihood from the LogR); and the second and third steps are repeated a hundred times or until convergence.

Using the major minor configuration of the data as described above, MEDICC2 was run with standard settings on 32 cores for patient TN1 and TN2 of the cohort. By looking at the final tree and the corresponding copy-number profiles, clades in the tree were manually assigned to the corresponding super- and subclone of the original publication. In order to recreate the minimum evolution trees from the original publication [41] we created phylogenies using the function *fastme*.*bal* from the R package ape [63] based on the pairwise Manhattan distance.

## Implementation and Availability

MEDICC2 is implemented in Python 3 and freely available under GPLv3 at https://bitbucket.org/schwarzlab/medicc2. It employs OpenFST and its Python wrapper pywrapfst [60] for manipulation of finite-state machines. Core algorithms are implemented in C++ as a Cython extension by linking to the OpenFST library. All data and code to reproduce the figures of this publication are present in the repository.

## Contributions

TLK and RFS designed the method; TLK, MP and RFS implemented the method; TLK and KH processed and analysed the PCAWG single sample copy-number data; TBKW, ECC and NT processed and analysed the PCAWG multi sample copy-number data; TLK and MT processed and analysed the single-cell data; DCM and NN provided the single-cell data; SL and TLK implemented the copy-number simulations; TLK, TBKW, KH, ECC, MT, CS, PVL, and RFS wrote the manuscript.

## Acknowledgements

R.F.S., T.K., and M.P. thank the Helmholtz Association (Germany) for support. T.K. was funded by the German Ministry for Education and Research as BIFOLD - Berlin Institute for the Foundations of Learning and Data (ref. 01IS18025A and ref 01IS18037A). T.K. kindly thanks Klaus-Robert Müller for support. T.B.K.W thanks the Foulkes Foundation for support. T.B.K.W. and C.S. were supported by a Royal Society Research Professorships Enhancement Award (RP/EA/180007), the Breast Cancer Research Foundation (BCRF), Marie Curie ITN Project PLOIDYNET (FP7-PEOPLE-2013, 607722). T.B.K.W., C.S., M.T. and P.V.L were supported by the Francis Crick Institute, which receives its core funding from Cancer Research UK (FC001169, FC001202), the UK Medical Research Council (FC001169, FC001202) and the Wellcome Trust (FC001169, FC001202). For the purpose of Open Access, the authors have applied a CC BY public copyright licence to any Author Accepted Manuscript version arising from this submission. C.S. is a Royal Society Napier Research Professor (RP150154). M.T. was supported as a postdoctoral researcher of the F.R.S.-FNRS. This project was enabled through access to the MRC eMedLab Medical Bioinformatics infrastructure, supported by the Medical Research Council (grant number MR/L016311/1). P.V.L. is a Winton Group Leader in recognition of the Winton Charitable Foundation’s support towards the establishment of The Francis Crick Institute. Computation has been performed on the HPC for Research cluster of the Berlin Institute of Health.

