## Supplemental Figures for "MEDICC2: whole-genome doubling aware copy-number phylogenies for cancer evolution"

[1]*Berlin Institute for Medical Systems Biology, Max Delbrück Center for Molecular Medicine in the Helmholtz Association (MDC), Robert-Rössle-Str. 10, 13125 Berlin, Germany.*
[2]*Department of Electrical Engineering & Computer Science, Technische Universität Berlin, Marchstr. 23, Berlin 10587, Germany.*
[3]*BIFOLD, Berlin Institute for the Foundations of Learning and Data, Berlin, Germany.*
[4]*Department of Biology, Humboldt University of Berlin, Unter den Linden 6, 10099 Berlin, Germany.*
[5]*Charité – Universitätsmedizin Berlin, corporate member of Freie Universität Berlin and Humboldt Universität zu Berlin, Department of Pediatrics, Division of Oncology and Hematology, Augustenburger Platz 1, 13353 Berlin, Germany.*
[6]*The Francis Crick Institute, London, UK.*
[7]*Department of Mathematics and Computer Science, Free University of Berlin, Germany.*
[8]*UCL Medical School, University College London, London, UK*
[9]*Department of Genetics, The University of Texas MD Anderson Cancer Center, Houston, TX, USA*
[10]*Cancer Research UK Lung Cancer Centre of Excellence, University College London Cancer Institute, London, UK.*
[11]*Department of Medical Oncology, University College London Hospitals, London, UK.*
[12]*German Cancer Consortium (DKTK), German Cancer Research Center (DKFZ), Heidelberg, Germany.*
[13]*Institute for Interdisciplinary Research, Université Libre de Bruxelles, Brussels, Belgium*
[*]*these authors contributed equally*
[@]*contact:*

### Supplementary Figures

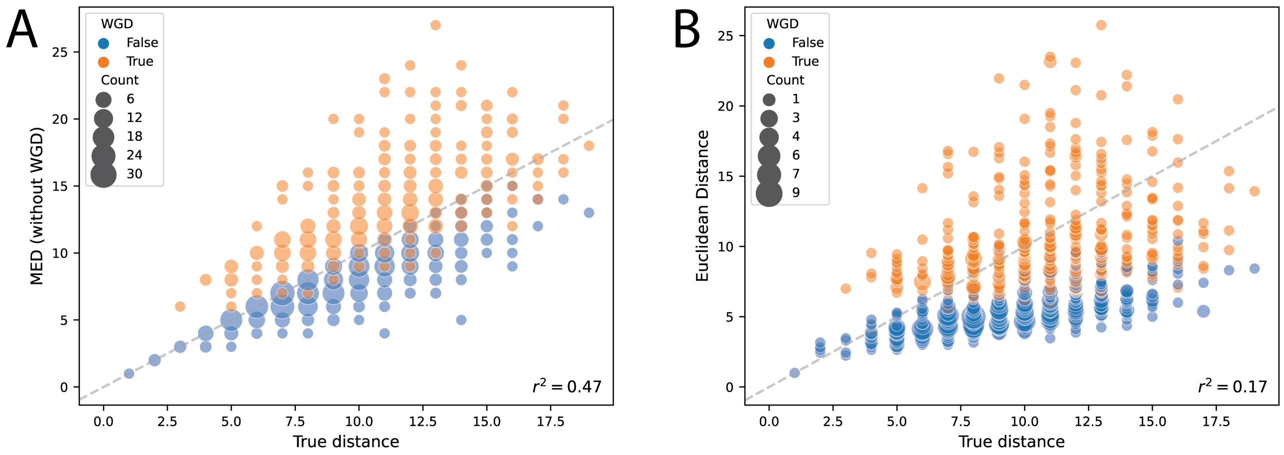


**Supplementary Figure 1: Model performance on simulated distances A)** MEDICC2 overestimates distances in sequences where WGD events have occurred if using the MED without taking WGD events into account. **B)** Euclidean distance is unable to recover the actual tree distance, especially if WGD events have occured.

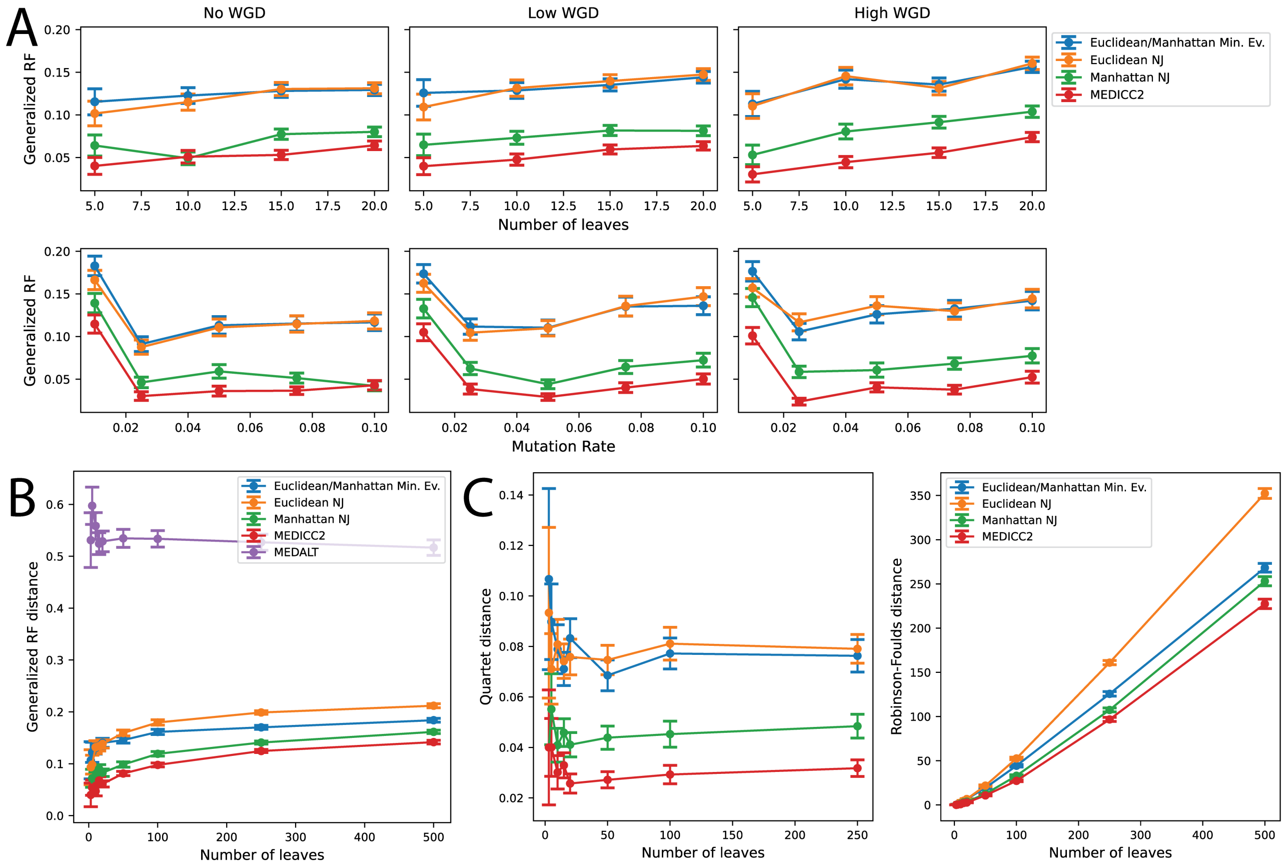


**Supplementary Figure 2: Model performance on simulated trees. A)** Results of the tree reconstruction for a range of intermediate tree sizes, reaction rates and WGD rates. **B)** Results for the phylogenetic tree reconstruction method *MEDALT* which creates minimum-spanning trees and therefore performs poorly on the simulated trees. **C)** MEDICC2 outperforms all other methods also for two other, widely-used tree distance measures; the Quartet distance and the Robinson-Foulds distance.

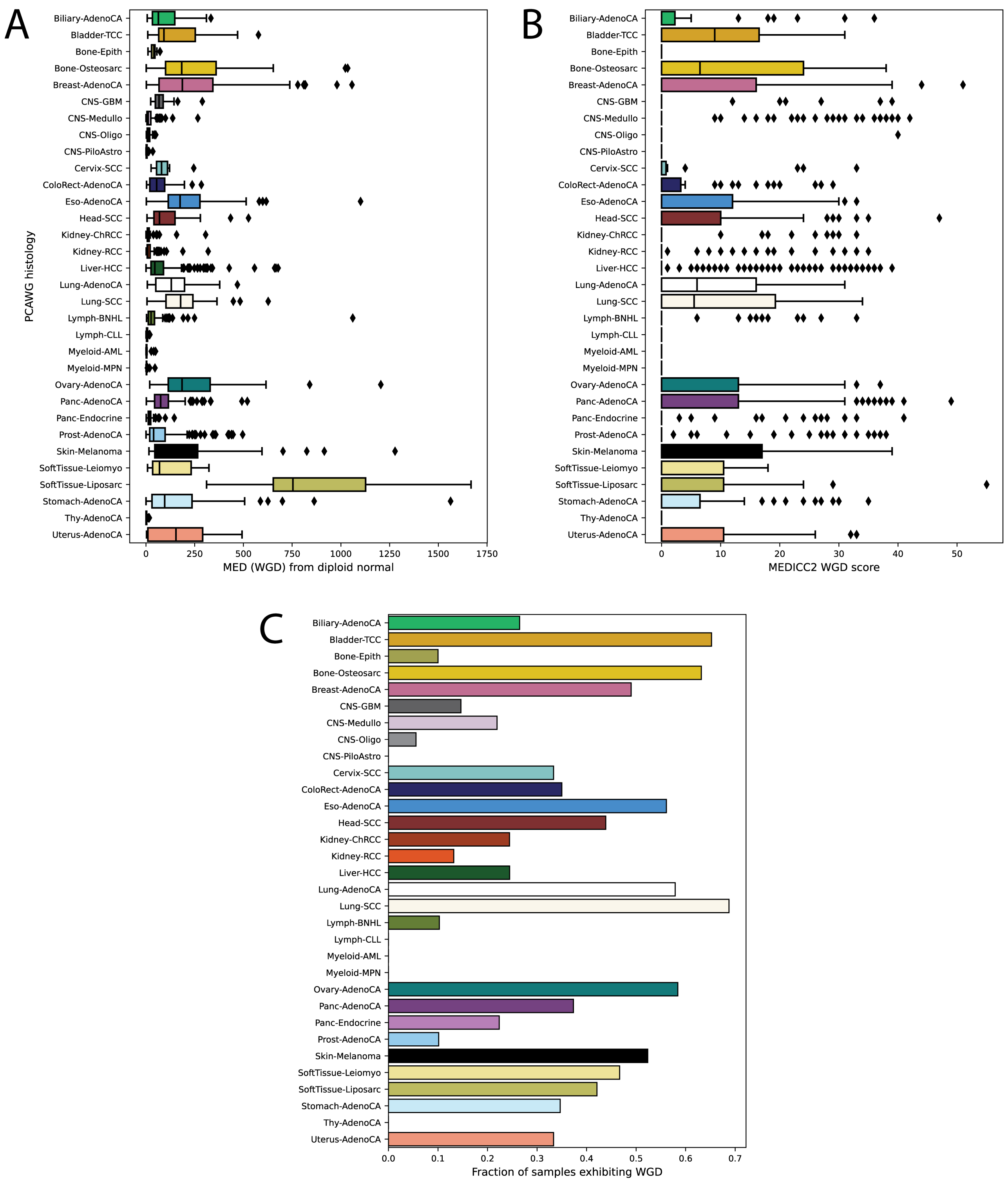


**Supplementary Figure 3:** **Overview of the PCAWG cohort. A) MEDICC2 MED-WGD for all PCAWG tumours** show differing levels of copy-number evolution. **B) MEDICC2 WGD score for all PCAWG tumours** and **C) Fraction of samples exhibiting WGD** shows varying degrees of WGD events across the cohort.

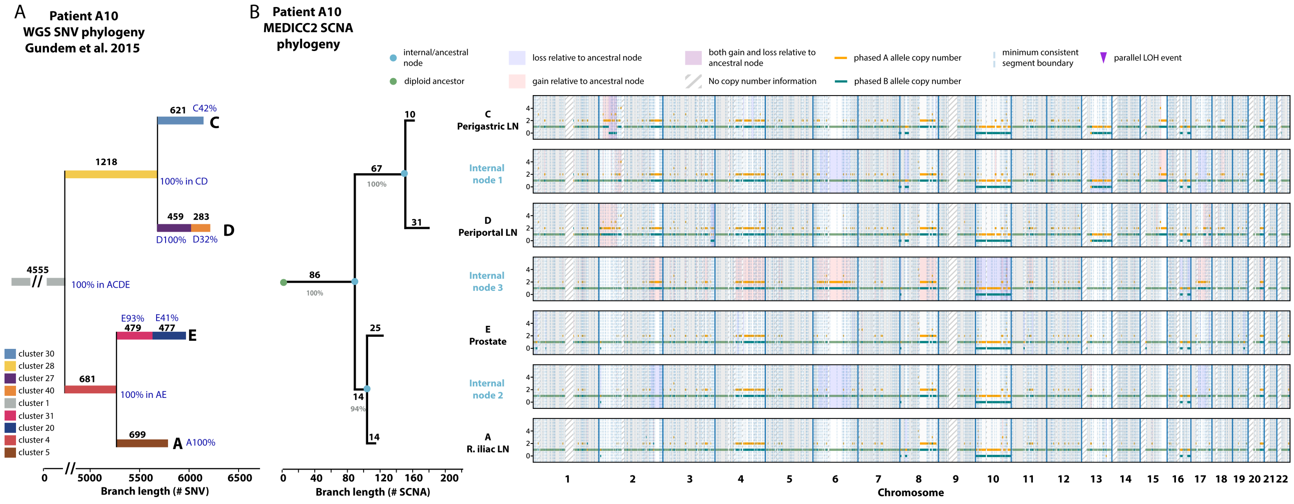


**Supplementary Figure 4:** Evolutionary history of tumour subclones from patient ”A10”.

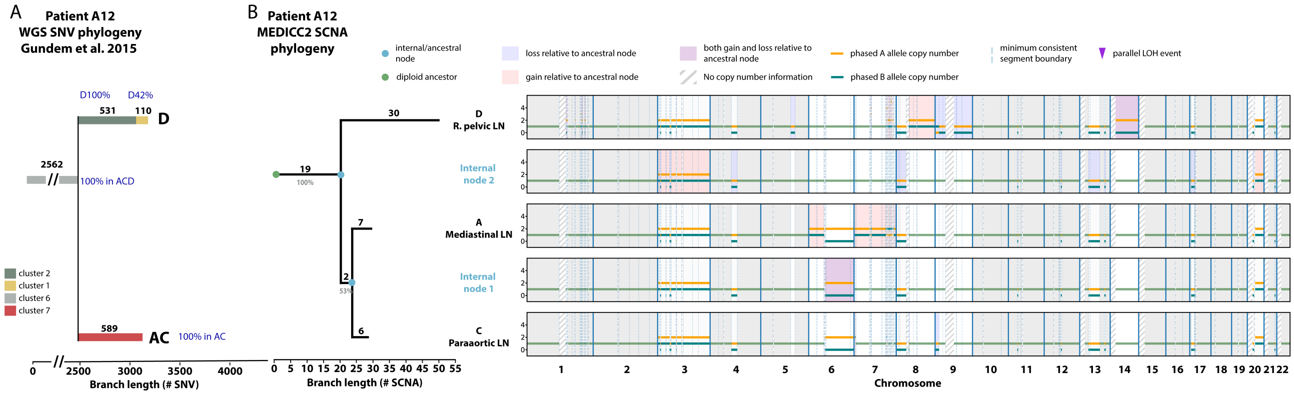


**Supplementary Figure 5:** Evolutionary history of tumour subclones from patient “A12”.

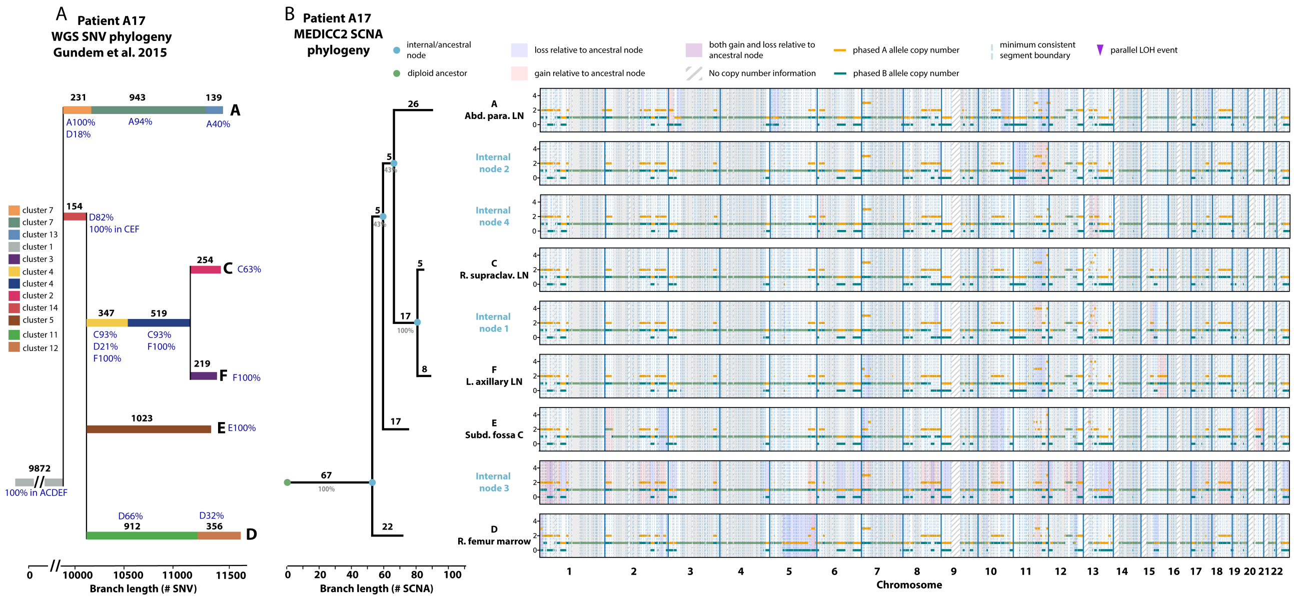


**Supplementary Figure 6:** Evolutionary history of tumour subclones from patient “A17”.

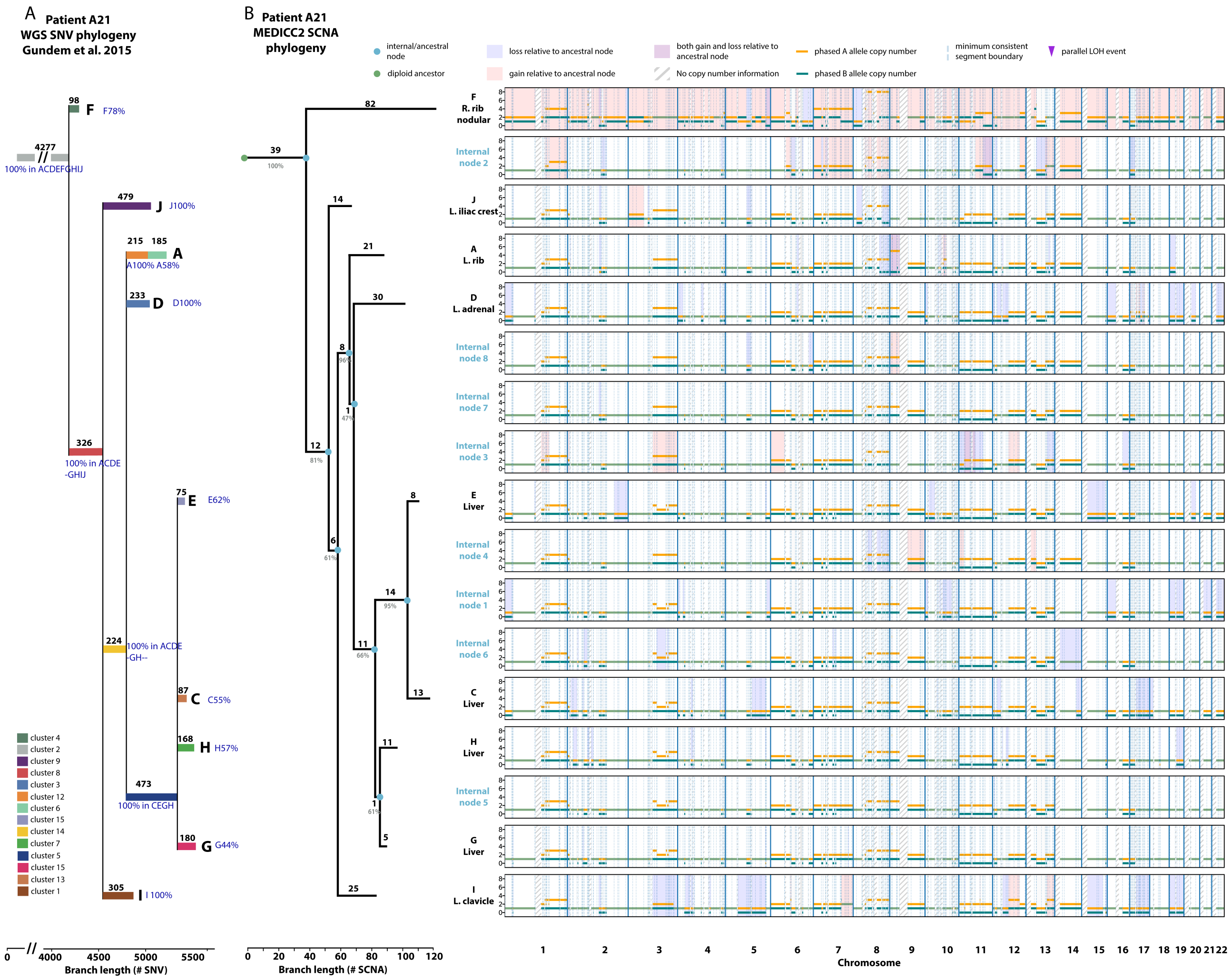


**Supplementary Figure 7:** Evolutionary history of tumour subclones from patient "A21".

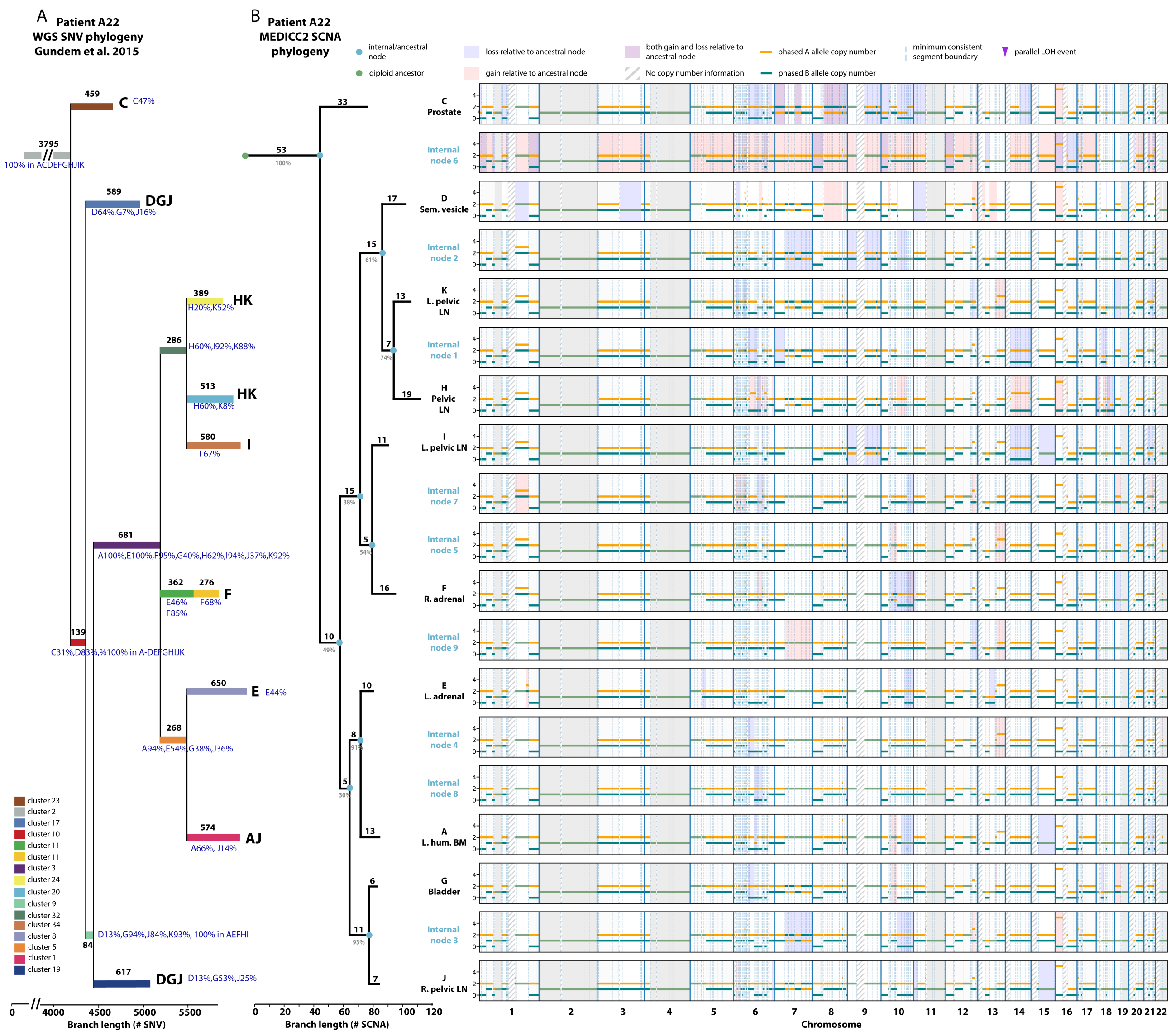


**Supplementary Figure 8:** Evolutionary history of tumour subclones from patient “A22”.

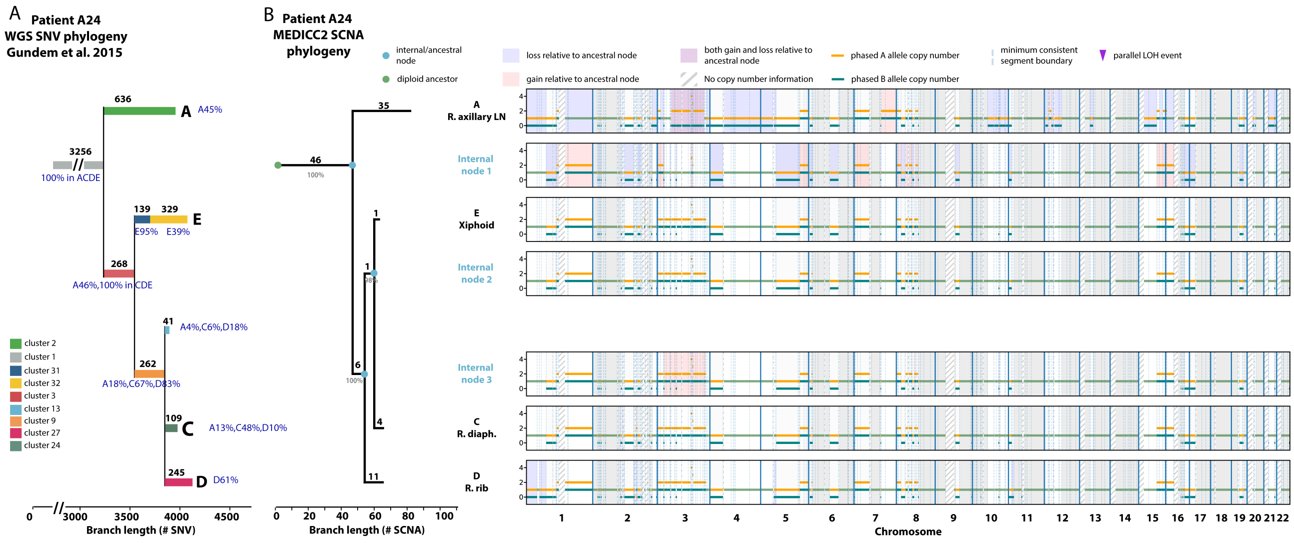


**Supplementary Figure 9:** Evolutionary history of tumour subclones from patient “A24”.

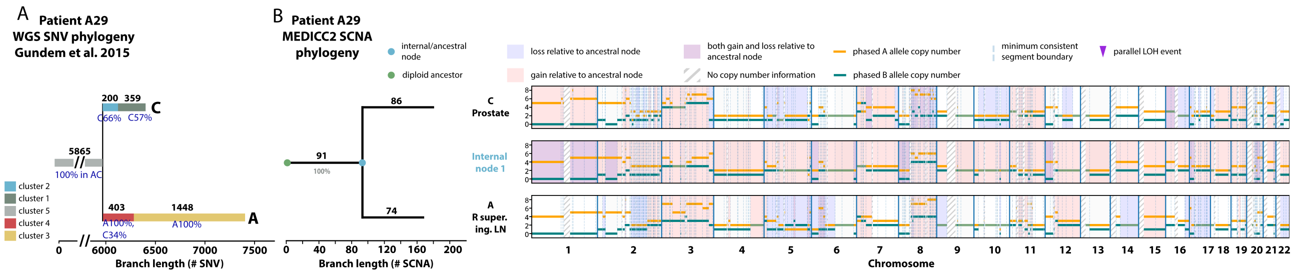


**Supplementary Figure 10:** Evolutionary history of tumour subclones from patient "A29".

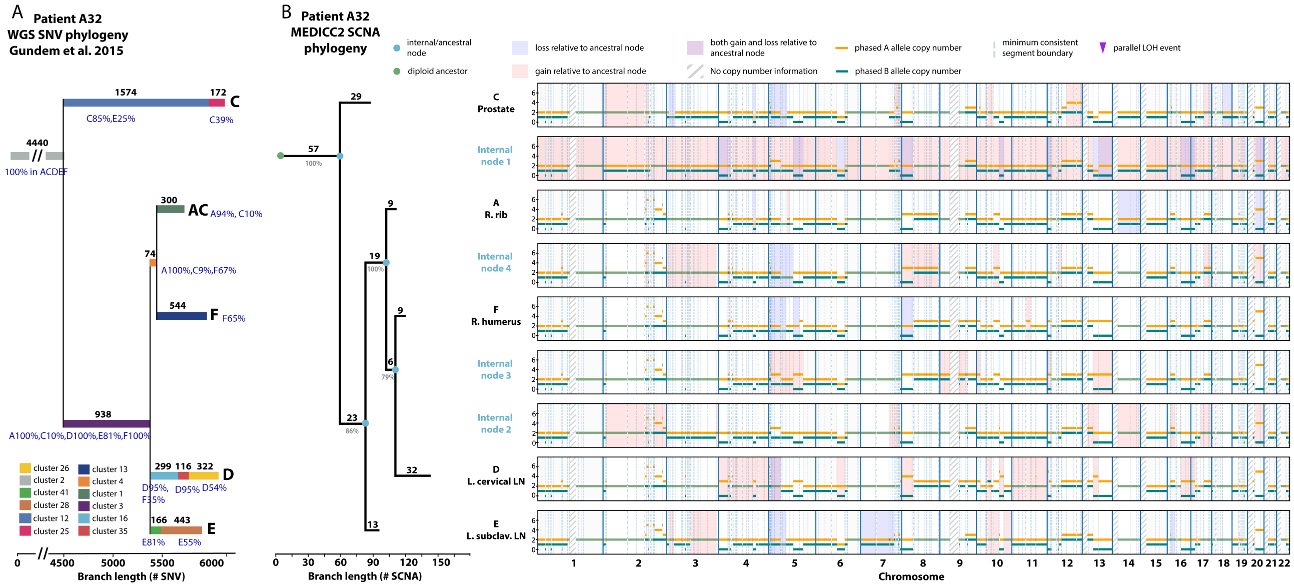


**Supplementary Figure 11:** Evolutionary history of tumour subclones from patient "A32".

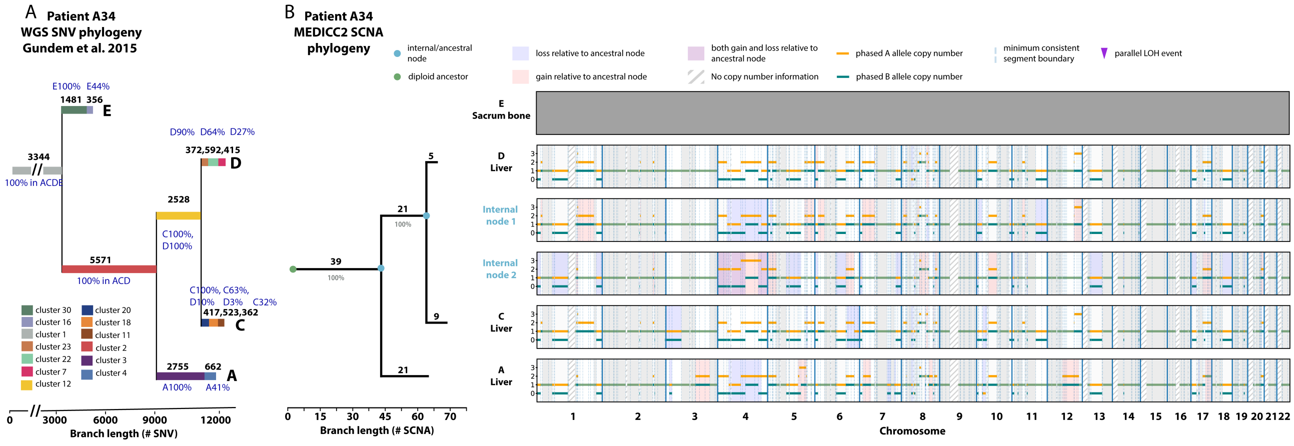


**Supplementary Figure 12:** Evolutionary history of tumour subclones from patient "A34".

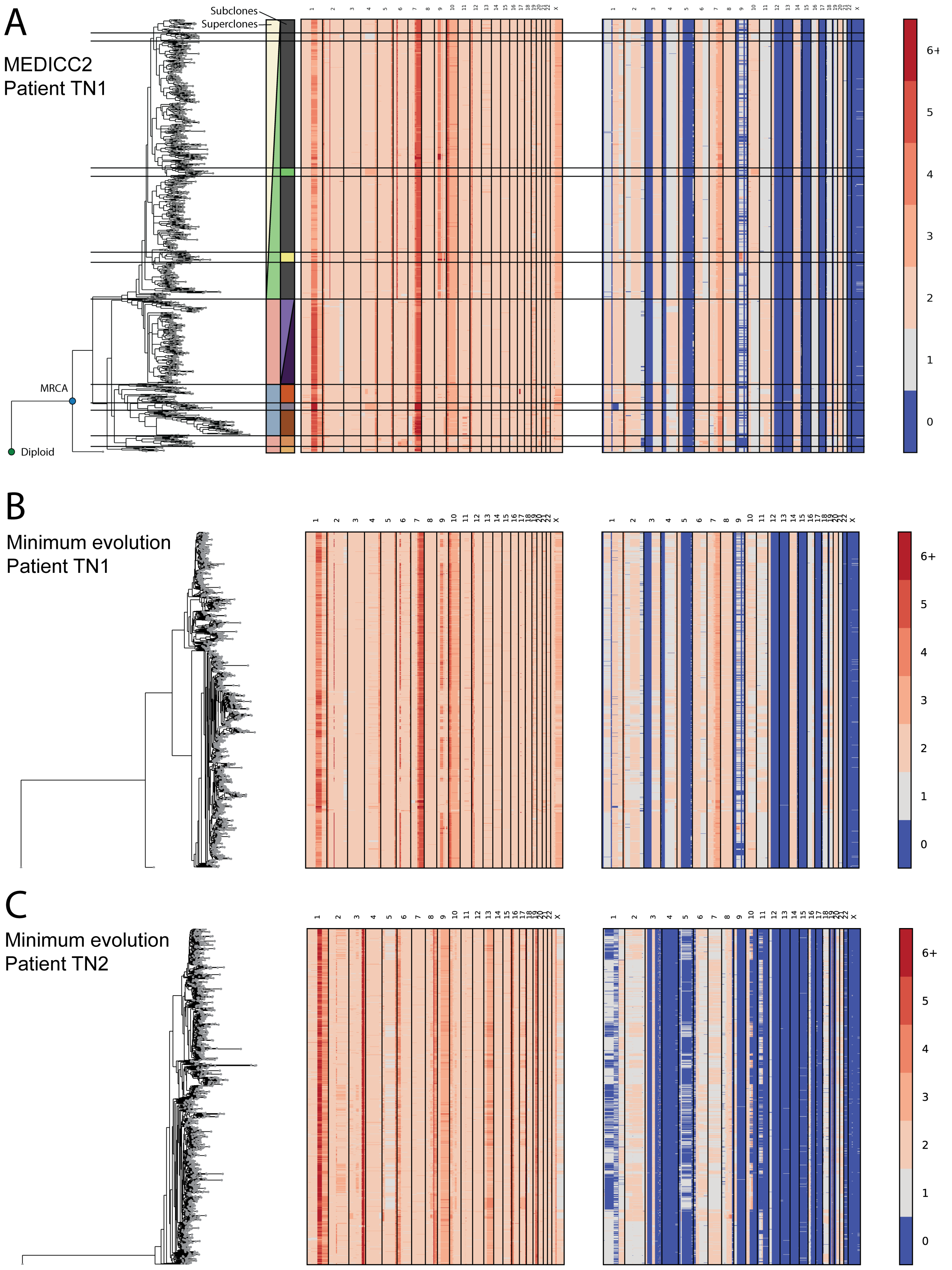


**Supplementary Figure 13:** **Single-cell cohort from Minussi et al. 2021** **A)** Inferred phylogeny and allele-specific copy-number profiles for patient TN1 from Minussi et al. 2021. Superclones and subclones are marked as in the original publication. **B)** and **C)** show the inferred phylogenies using the manhattan-distance based minimum evolution tree as described in the original publication. Unlike the MEDICC2 trees, the Manhattan trees do not recover the super- and subclone structure.
